# A Cationic Amphiphilic Drug (CAD) Defense System in the Nematode *Caenorhabditis elegans*

**DOI:** 10.64898/2026.01.03.697488

**Authors:** Levon Tokmakjian, Duhyun Han, Kateryna Sihuta, Aanchal Aggarwal, Somayeh Pirhadi, Yao Wang, Andrew R. Burns, Brittany Cooke, Siyue Ren, Carolyn L. Cummins, Justin Nodwell, David R. Koes, Peter J. Roy

**Affiliations:** Department of Pharmacology and Toxicology, University of Toronto, Toronto, ON, M5S 1A8, Canada; The Donnelly Centre for Cellular and Biomolecular Research, University of Toronto, Toronto, ON, M5S 3E1, Canada; Department of Molecular Genetics, University of Toronto, Toronto, ON, M5S 1A8, Canada; Department of Computational and Systems Biology, University of Pittsburgh, 4200 Fifth Ave, Pittsburgh, Pennsylvania 15260, United States; Department of Pharmaceutical Sciences, Leslie Dan Faculty of Pharmacy, University of Toronto, Toronto, Ontario, Canada; Department of Biochemistry, University of Toronto, 1 King’s College Circle, Toronto, ON, M5S 1A8, Canada

## Abstract

Cationic Amphiphilic Drugs (CADs) severely disrupt lysosomal function, which leads to a cellular pathology characterized by excess phospholipids called phospholipidosis. Through a forward genetic screen and mining of published datasets, we discovered that CADs induce the expression of the CYP-35B family of cytochrome P450s and the PGP-13 p-glycoprotein pump via the nuclear receptors NHR-70 and NHR-107 in the nematode *C. elegans*. A *pgp-13* fluorescent reporter revealed hundreds of human drugs that upregulate the CAD defense system *in vivo*. Chemoinformatic analyses indicate that the *pgp-13* reporter may be useful in identifying CADs that have pathogenic potential in humans. Mutant analyses coupled to metabolomics and structural modeling show that the CYP-35Bs are necessary and sufficient for CAD metabolism, and that CYP-35B2 D311 is key in mediating electrostatic interactions with the positively charged CADs. We also show that CAD metabolites are effluxed via PGP-13 acting partially redundantly with PGP-14 and that an intact defense system is necessary to resist CAD-induced pathology. Finally, we demonstrate that bacteria that likely cohabitate with *C. elegans* in nature trigger the CAD defense system, providing a plausible explanation for why a pathway that protects against anthropogenic small molecules exists in nematodes.

## Introduction

Cationic Amphiphilic Drugs (CADs) can severely disrupt lysosomal function and lead to a cellular pathology called drug-induced phospholipidosis (PLD) (Breiden AND Sandhoff 2019). CADs are small lipophilic molecules that typically have a secondary or tertiary amine with a pKa around 8 (Halliwell 1997). At physiological pH, a good fraction of the CAD molecules is unprotonated and can pass through membranes. In acidic conditions, a greater fraction of the CAD molecules is positively charged and become trapped in acidic organelles like the lysosome and accumulate there to high concentrations (Tummino *et al*. 2021). The protonated CADs neutralize the negatively charged membranes of the acidic organelles (Breiden AND Sandhoff 2019). Consequently, the phospholipid-degrading enzymes that normally interact with the lysosome via its negative membrane charge can no longer do so in the presence of CADs (Abe AND Shayman 2009). In this way, CADs drive lipid accumulation (aka PLD) as an unintended side-effect.

CADs are problematic for several reasons. First, CADs are more likely to be associated with drug-induced liver injury (DILI) compared to non-CADs (Chen *et al*. 2013). DILI can lead to liver transplantation and death for many of those afflicted (Chalasani *et al*. 2015; Dirven *et al*. 2021). Second, most PLD-inducing CADs have physicochemical properties (i.e., a hydrophobic backbone coupled to a positively charged atom) in common with blockers of the human ether-a-go-go related gene product hERG (KCNH2 or Kv11.1) (Creanza *et al*. 2021). Indeed, 77% of PLD-inducing CADs have been shown to block hERG activity (Sun *et al*. 2013). hERG is a voltage-gated potassium channel that is key in maintaining cardiac rhythm and whose disruption can lead to sudden cardiac death (Vandenberg *et al*. 2012). Given their associated DILI and cardiac pathologies, cationic amphiphilic molecules are considered high risk in medicinal chemistry pipelines (Pelletier *et al*. 2007; Vandenberg *et al*. 2012). Despite this, roughly 5% of all small molecule drugs used in the clinic are PLD-inducing CADs (Schieferdecker *et al*. 2022).

A master drug clearance pathway in mammals consists of the PXR nuclear receptor, which when bound by a variety of compounds, will drive the expression of multiple drug-metabolizing cytochrome P450s and the drug efflux pump ABCB1 (aka MDR1, aka Pgp) (Blumberg *et al*. 1998; Geick *et al*. 2001; Takeshita *et al*. 2002). However, known drug detoxification pathways are only modestly effective when responding to CADs. For example, although the canonical CAD amiodarone induces ABCB1 expression, the drug inhibits the pump’s efflux activity (Cermanova *et al*. 2010; Hodges *et al*. 2011). Similarly, while multiple cytochrome P450s can metabolize amiodarone, the resulting metabolites inhibit these same P450s (McDonald *et al*. 2015). Another classic CAD is chlorpromazine, which activates the PXR drug response (Faucette *et al*. 2007). While the P450s CYP1A2, CYP2D6, and CYP3A4 metabolize chlorpromazine (Boyd-Kimball *et al*. 2019), chlorpromazine itself inhibits ABCB1’s activity (Hodges *et al*. 2011).

Here, we have discovered a coordinated CAD defense system consisting of the nuclear receptors NHR-70 and NHR-107, the CYP-35B P450 family, and the PGP-13 and PGP-14 efflux pumps using the nematode worm *C. elegans*. Through our previous small molecule screens (Burns *et al*. 2015), we discovered one class of small molecules (not CADs) that kills the nematode *C. elegans* by forming crystals exclusively in the pharynx cuticle (Kamal *et al*. 2019; Kamal *et al*. 2022; Kamal *et al*. 2023; Kamal *et al*. 2024). These small molecule crystals kill young nematodes because they pierce the underlying epithelium of the pharynx cuticle and grow to occlude the lumen (Kamal *et al*. 2023).

Forward genetic screens revealed that the p-glycoprotein (PGP) pump PGP-14 is key in facilitating small molecule crystal formation; without it, crystals do not form (Kamal *et al*. 2023). We found PGP-14 to be expressed exclusively in the pharynx epithelium that abuts the pharynx cuticle (Kamal *et al*. 2023). There, PGP-14 is needed for polar lipid accumulation within the cuticle, suggesting that PGP-14 exports phospholipids into the cuticle where they may protect the animal from water loss and xenobiotic attack (Kamal *et al*. 2019; Kamal *et al*. 2023). We speculate that the lipids within the pharynx cuticle act as a sink for the accumulation of select hydrophobic small molecules where they precipitate out of solution and kill the animal (Kamal *et al*. 2023; Kamal *et al*. 2024).

PGP-14 is most homologous to the human PGPs ABCB4, a pump that exports phospholipids from hepatocytes to form bile (Smit *et al*. 1993; Prescher *et al*. 2019), and the aforementioned ABCB1 multidrug efflux pump (Alam *et al*. 2019; Nosol *et al*. 2020; Kamal *et al*. 2023). Without functional ABCB4, phospholipids are not present to promote bile acid entry within the hepatocytic duct, leading to hepatocytic damage that can result in a childhood liver disease called progressive familial intrahepatic cholestasis 3 (PFIC3) (Park *et al*. 2016). There is a clear analogy to PGP-14’s role in establishing the lipid barrier within the *C. elegans* pharynx cuticle and ABCB4’s role in phospholipid export from hepatocytes.

Given PGP-14’s analogous role and homology to ABCB4, we were curious to explore the potential utility of the *pgp-14(0)* null mutant as a model for ABCB4 deficiency in humans. Towards this end, we screened the Spectrum library of 2560 drugs and natural products for those that suppress *pgp-14(0)*. All resulting suppressors are structurally diverse CADs. Through a forward genetic screen, we discovered that CADs suppress *pgp-14(0)* because they upregulate an ABCB1 homolog called PGP-13 that acts redundantly with PGP-14. In addition to PGP-13, CADs upregulate multiple cytochrome P450s of the CYP-35B family, via the nuclear receptor transcription factors NHR-70 and NHR-107.

Using mass-spectrometry, we show that the P450s metabolize CADs into multiple products, some of which are effluxed by PGP-13 acting either alone or redundantly with PGP-14. In this way, we have discovered a novel coordinated CAD defense pathway, without which the animals become more sensitive to the pathological effects of CADs.

To better understand the diversity of molecules that engage the CAD defense system we built a fluorescent reporter of *pgp-13*. We used this reporter to screen the Spectrum small molecule library at a single concentration and identified 293 structurally diverse inducers, 78% of which have either secondary or tertiary amines, indicating that *C. elegans* has a robust and plastic CAD detection system. A published chemoinformatic predictor of PLD**-**inducing CADs (Pelletier *et al*. 2007) predicts that 73% of our 293 hits are PLD-inducing CADs. The same chemoinformatic tool predicts the same fraction of PLD-inducing CADs from a set of 112 gold standard PLD-inducing CADs that we assembled from the literature. Hence, our *in vivo* CAD defence reporter is at least as good as a computational predictor of CAD-induced pathologies in humans and provides a facile model for testing predictions.

Finally, we asked whether bacteria that likely cohabitate with *C. elegans* in nature produce products that engage the CAD-defense system. We screened 65 *Streptomyces* strains, the genus of which are highly abundant within the natural environment of *C. elegans* (Kampfer 2006; Samuel *et al*. 2016) and found two strains that robustly upregulate the CAD defense reporter in a nuclear receptor-dependent manner. These observations suggest that the CAD defense system exists in nematodes to defend against natural product CADs that have the potential to harm the worm.

## Results

### A Screen for Small Molecules that Rescue *pgp-14(0)* Yields Multiple CADs

To determine whether the phenotypes associated with the *pgp-14(0)* null mutant could be suppressed by small molecules, we screened 2560 clinically approved drugs and natural products of the Spectrum library (Microsource Inc) for those that can rescue *pgp-14(0)*’s resistance to the lethal crystal-forming wact-190 (3-(4-(benzyloxy)phenyl)-5-(pyridin-4-yl)-1,2,4-oxadiazole) (Figure 1A) (Kamal *et al*. 2023). We focused on 14 reproducible hits (Figure 1B; Source Data File 1-tab 1, column ‘Z’). These hits rescued *pgp-14(0)*’s resistance to both the death/arrest phenotype induced by crystal-forming molecules (Figure 1C) and the ability of the molecules to form crystals (Figure 1D). In the wildtype, crystal-forming molecules form obvious crystals over 24 hours. In the *pgp-14(0)* mutant, crystals rarely if ever form. *pgp-14(0)* mutants grown in the rescuing drugs take about two days to form obvious crystals in the arrested animals. Hence, the 14 rescuing compounds have slower temporal dynamics compared to wild type PGP-14.

**Figure 1.**
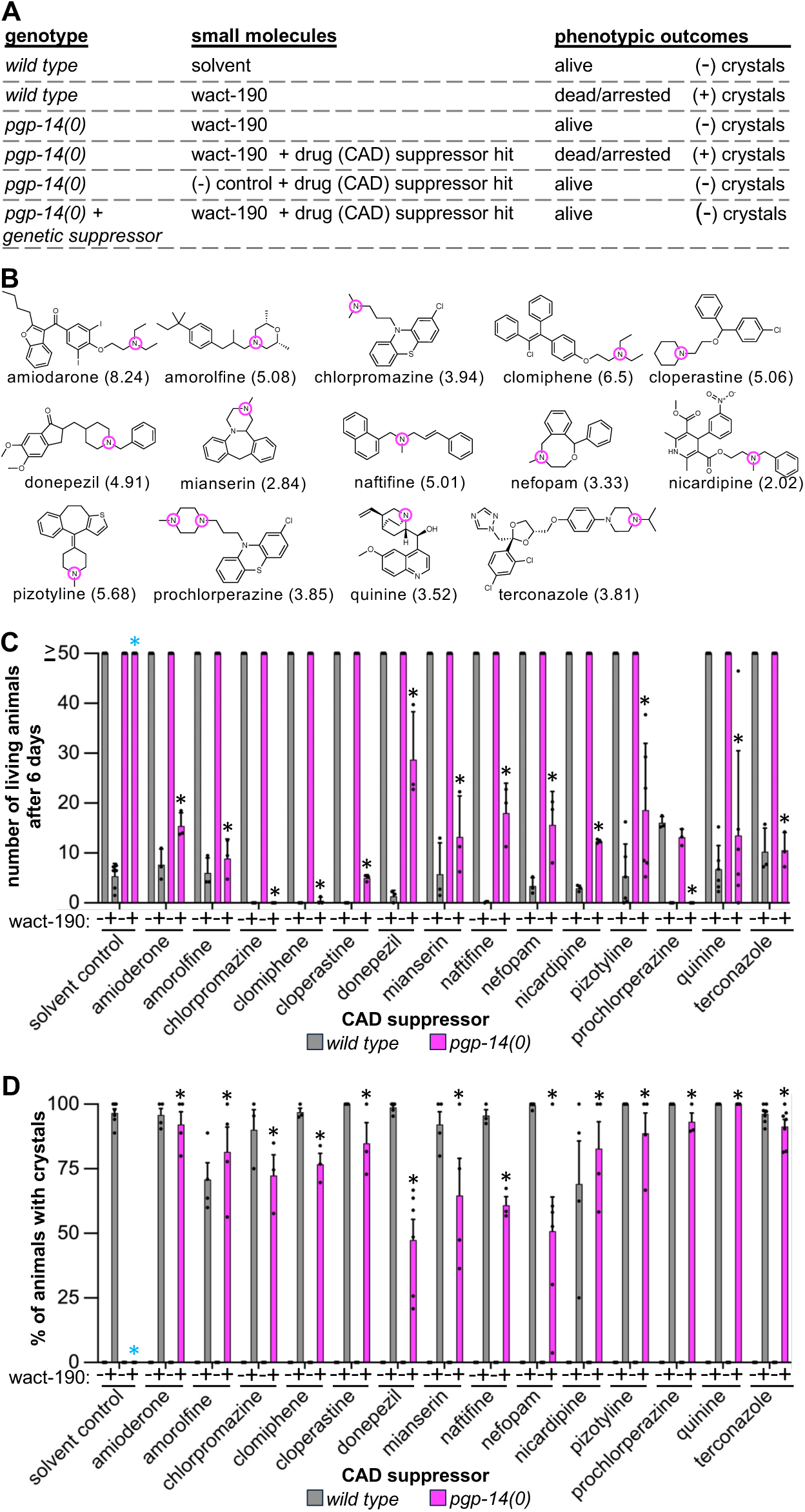
Drug suppressors of *pgp-14(0)*. **A.** A schematic illustrating the logic of the drug suppressor screen of *pgp-14(0)*. 30 μM of drug and 15 μM of wact-190 were used in the screen. **B.** Silicos-IT Log P estimations are indicated in brackets. **C.** Six-day viability assays illustrating how select CADs, tested at 30 μM, resensitize the *pgp-14(0)* mutant *ok2660* to the lethality of 30 μM wact-190. Three independent repeats were performed (N>3); an upper ceiling of 50 animals per sample were counted. An asterisks indicates *p*<0.001 in a Student’s 2-tailed T-test, calculated only for the *pgp-14(0) +* wact-190 condition relative to the *pgp-14(0) +* solvent-only control (indicated with a blue asterisks). **D.** 30 μM of wact-190 and 30 μM of the indicated drug were used. At least 3 independent repeats were performed (N≥3), with at least 5 worms counted per sample (n≥5). An asterisks indicates *p*<0.05 in a Student’s 1-tailed T-test, calculated only for the *pgp-14(0)* background relative to the solvent-only control (indicated with a blue asterisks). For both graphs, standard error of the mean is shown. See Source Data File 1 for details.

The 14 rescuing compounds are: i) drugs used in the clinic for diverse indications, ii) structurally diverse, iii) tertiary amine-containing molecules (and therefore positively charged at pH ≤ 7), and iv) hydrophobic molecules as measured by a computationally predicted hydrophobicity score (cLogP) greater than +2 (Figure 1B; Source Data File 1-tab 1). Hence, these 14 compounds are considered cationic amphiphilic drugs (CADs) (Halliwell 1997). The Spectrum library has a total of 336 tertiary amine-containing molecules (Source Data File 1-tab 1, column ‘FZ’). A hypergeometric *p*-value calculation indicates that the chances of all 14 hits being tertiary amine-containing molecules by random chance alone is very low (*p*≤4E-13).

### CADs Compensate for the Loss of PGP-14 Through the Upregulation of PGP-13

To understand how the CADs rescue *pgp-14(0)*, we carried out a forward genetic screen for mutants that suppress the ability of CADs to rescue *pgp-14(0)*. We grew randomly EMS-mutagenized *pgp-14(0)* mutants in a solution containing a crystal-forming compound (wact-190) and a CAD (terconazole). Normally, the CAD resensitizes the *pgp-14(0)* mutant to the lethality conferred by wact-190 crystal formation. We simply selected for mutants that could live under these conditions (see the bottom line of Figure 1A).

The results of the screen yielded multiple alleles of *nhr-70*, *nhr-107*, *pgp-13,* and *tat-3* (Figure 2A; Source Data File 1-tab 4). For each of the four genes, there are alleles that are highly likely to represent loss-of-function alleles, yielding either early stops, deletions, or splicing defects (Figure 2A; Supplemental Figure 1). We do not further discuss *tat-3* here. NHR-70 and NHR-107 are distantly related to the human HNF4 family of nuclear receptors and do not have clear one-to-one human orthologs (Robinson-Rechavi *et al*. 2005). In humans, the HNF4s are primarily expressed in endodermal epithelia, including the liver and intestine (Sasaki *et al*. 2018; Vemuri *et al*. 2023) and are closely related to the xenobiotic-responsive PXR and CAR nuclear receptors (Bookout *et al*. 2006). In blast searches with the PGP-13 query, human ABCB1 is the best human match. Of note, the *pgp-13* gene is situated immediately next to *pgp-14* in the genome, which is remarkably similar to the physical relationship between ABCB1 and ABCB4 in humans (Supplemental Figure 2).

**Figure 2.**
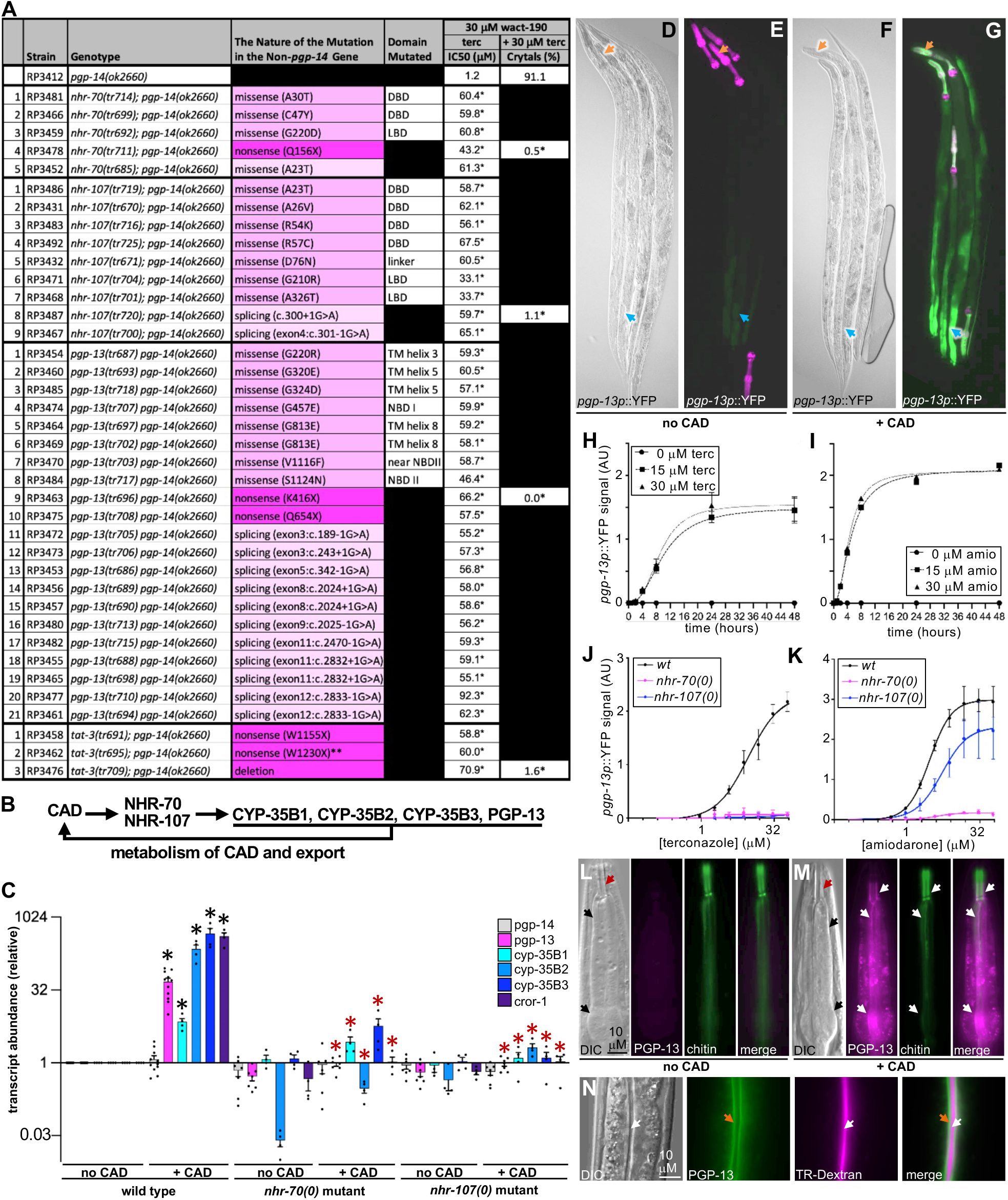
Components of a Putative CAD-Defense System. **A.** The mutant genes that suppress terconazole’s ability to resensitize *pgp-14(0)* to wact-190-crystal-induced death. The double asterisk indicates that *tr695* also has splicing mutation in exon 3 of uncertain consequence. The IC50 of terconazole’s ability to resensitize the different mutants to 30 micromolar wact-190 is shown, demonstrating the mutants’ resistance to terconazole. Wact-190 crystal formation in select mutants is also shown. A single asterisk indicates *p*<0.001 with 2-way ANOVA using the 30 μM concentrations. **B.** A model of the CAD-defense system. CADs engage nuclear receptor transcription factors (directly or indirectly), which promote the expression of cytochrome P450s (that may metabolically modify CAD structure, potentially altering its amphiphilic nature and making it a better substrate for export by PGPs) and PGP-13, which may act in concert with PGP-14 to export the CAD out of the cell. **C.** qRT-PCR analyses of the expression of putative CAD-defense factors. The CAD used is 30 μM terconazole. N≥4 independent repeats. The black asterisk indicates *p*<0.05 relative to the no-CAD solvent-only control in wild type animals. The red asterisk indicates a significant difference (*p*<0.05) between the mutants + CAD relative to the wild type + CAD. **D & E.** Four RP3507 *trIs114* [*pgp-13p::YFP; p382(myo-2p::mCherry*)] animals incubated in 1% DMSO solvent for 24 hours. **F & G.** Four RP3507 *trIs114* (*pgp-13p*::YFP; p382(*myo-2p*::mCherry)) animals incubated in 30 μM terconazole for 24 hours. Differential interference contrast (DIC) images are shown in D and F, darkfield composites of the RFP channel (fuchsia) and YFP channel (green) are shown in E and G. The blue arrow highlights the intestine of one animal; the orange arrow highlights the procorpus of the pharynx of the same animal. **H & I.** Temporal response of the *trIs114 pgp-13p*::YFP reporter to the indicated CAD in a wild type (RP3507) background. N=3 independent repeats with 72 technical repeats per trial (n=72). **J & K.** Dose-response of the *trIs114 pgp-13p*::YFP reporter to the indicated CAD (30 μM) in a *wild type* (RP3507), *nhr-70(tm1697)* (RP3518) and *nhr-107(tm4512)* (RP3516) background. N=3 independent repeats, n=12 per trial. **L & M.** RP3561 *trIs115* [CRISPR mNeonGreen KI N-terminal fusion with PGP-13) in *pgp-14(ok2660)]* incubated in either solvent control (L) or 30 μM of the CAD terconazole for 48 hours and then stained for chitin with calcofluor white (green). The mNeonGreen::PGP-13 is shown in fuchsia. Only the head of the animal is shown with the buccal cavity highlighted with a rust arrow and the procorpus of the pharynx highlighted with black arrows in the DIC images. The localization of the mNeonGreen::PGP-13 to the apical membrane of the pharynx myoepithelium is indicated with white arrows. **N.** Localization of mNeonGreen::PGP-13 (green) to the apical membrane of the intestine. DIC (left) with a white arrow indicating the lumen. mNeonGreen::PGP-13 (orange arrow) is localized to the apical membrane, while Texas Red-Dextran (white arrow) is restricted to the intestinal lumen. See Source Data File 1 for details.

The screen results suggested that CADs directly or indirectly activate the NHR-70 and NHR-107 nuclear receptors to transcriptionally up-regulate PGP-13, the closest paralog of PGP-14 (Sheps *et al*. 2004; Kamal *et al*. 2023), to restore the *pgp-14(0)* mutant’s sensitivity to wact-190-mediated lethality (Figure 2B). We tested this model in two ways. First, we used qRT-PCR to examine *pgp-13* mRNA expression in response to the CAD terconazole in the background of the nuclear receptor mutants. We observed a 47-fold induction of *pgp-13* expression in response to the CAD terconazole (*p*=1.7E-05) and find that this upregulation is dependent on functional NHR-70 and NHR-107 (*p*=4.1E-04 for both) (Figure 2C). Second, we asked whether *pgp-13*, *nhr-70*, and *nhr-107* were necessary for the CAD’s ability to restore *pgp-14(0*)’s sensitivity to wact-190 crystal formation and found that they were (Supplemental Figure 3).

We explored the spatiotemporal expression pattern of *pgp-13* with a transcriptional YFP reporter, which revealed expression in the pharynx myo-epithelium and the intestine in a CAD and nuclear receptor-dependent manner (Figures 2D-2K). Expression is evident (*p*<0.05) within 2 hours of CAD exposure and peaks approximately 24 hours after initial exposure (Figure 2H and 2I; Source Data 1-tab 9). Intriguingly, one CAD’s ability to induce *pgp-13* expression (terconazole) is dependent on both NHR-70 and NHR-107, while another (amiodarone) is dependent on only NHR-70 (Figures 2J and 2K). We expand on these observations below. Upon CAD induction, a functional translational fusion reporter of PGP-13 (Supplemental Figure 4) shows subcellular localization to the apical membrane of the pharynx myoepithelium that is highly similar to that of PGP-14 (Kamal *et al*. 2023) (Figure 2L and 2M). The PGP-13 fusion protein also localizes to the apical membrane of the intestine (Figure 2N and 2O). Given the paralogy of PGP-13 and PGP-14 (Kamal *et al*. 2023) and the spatial overlap of their expression pattern, we infer that CAD-induced PGP-13 expression fortuitously compensates for the loss of *pgp-14* and accounts for why CADs were identified as *pgp-14(0)* suppressors in our initial screen.

### *pgp-13* is a Robust *in vivo* Marker of Diverse Biologically-Accessible CADs

We were curious to know what other small molecule structures might upregulate *pgp-13* expression in a nuclear receptor-dependant manner. We therefore screened 2400 compounds from the Spectrum library at a final concentration of 30 μM and identified 293 molecules that induce expression from the *pgp-13p*::YFP reporter (Source Data file 1-tab 1- column ‘J’). With each molecule reduced to its core scaffold (i.e., removing everything but rings, connectors and carbon chains), there are 195 distinct structural scaffolds, indicating that the hits are structurally diverse (Figure 3A). We evaluated our hits using two approaches. First, we compared our hits to a list of 112 molecules known to induce phospholipidosis in humans that we compiled from the literature and are also present in the Spectrum library (Morelli *et al*. 2006; Pelletier *et al*. 2007; Shahane *et al*. 2014; Bauch *et al*. 2015; Hinkovska-Galcheva *et al*. 2021) (Source Data file 1-tab 1 columns HU-IB). We refer to these molecules as gold standards. The *pgp-13*-inducers capture 68% of the gold standards, which is 5.6-fold more than expected by random chance (*p*=2E-46) (Figure 3B). To understand why 32% of the gold standards failed to induce *pgp-13* expression, we compared the physicochemical features of the gold standards that induce *pgp-13* expression compared to those that do not. We find that the latter are less likely to cross biological barriers (*p*<1E-5); they are significantly more polar (as measured by hydrogen bond donors and acceptors and total polar surface area), hydrophilic (as measured by LogP) and less likely to cross the skin and blood-brain barriers of humans (as measured by LogKp and a published predictive formula (Daina *et al*. 2017; Shaker *et al*. 2023), respectively) (Figure 3C). We surmise that most of the gold standards that fail to induce *pgp-13* expression would do so if assayed at higher concentrations, which would allow them to circumvent biological barriers. Consistent with this interpretation, many of the gold standards that failed to induce *pgp-13* expression have EC50 concentrations for PLD-induction in human cells that far exceed the concentration at which we screened in worms (30 μM) (e.g., amantadine, memantine, spectinomycin (Morelli *et al*. 2006); Source Data file 1-tab 1-column ‘HV’)).

**Figure 3.**
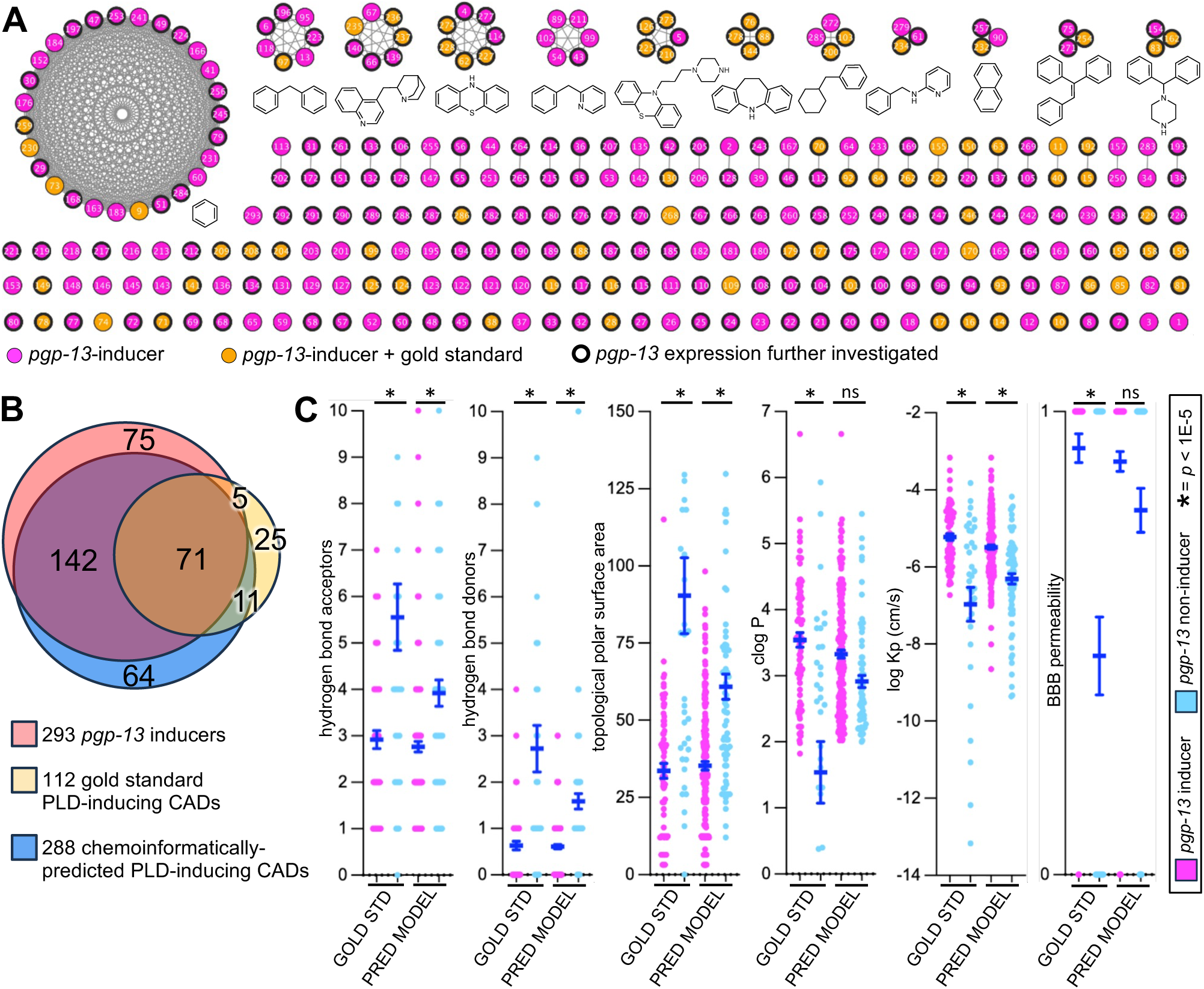
A Screen of 2400 Spectrum Small Molecules Reveals 293 Inducers of *pgp-13* expression. **A.** A structural similarity map of the 293 hits reveals 195 unique core scaffolds. Each of the nodes represents one of the 293 molecules reduced to its core scaffold (rings connected by chains). If two molecules within the collection of 293 molecules share the same scaffold, they are connected with a line. Gold-coloured nodes represent gold standard PLD-inducing molecules. Bold outlines are the 195 molecules further investigated in Figure 4. For clusters containing 3 or more molecules, the core scaffold is shown below, most of which have tertiary amines that are not part of the core scaffold. The identity of each molecule is represented by a number (see Source Data File 1-tab 1-column J to decode). **B.** Venn diagram showing the overlap between the *pgp-13*-inducing hits, the gold standards, and the predicted PLD-inducing CADs within the Spectrum library’s 2560 small molecules. **C.** Graphs of the select physicochemical properties (from SwissADME) of the molecules within the 112 gold standard (GOLD STD) set and the 288 predicted PLD-inducing CAD set (PRED MODEL) parsed on whether the molecule induces *pgp-13* expression or not. The physicochemical properties of all molecules are presented in Source Data File 1-tab 1. MLogP was used for LogP; LogKp is a measure of skin permeability (the higher the number the higher the permeability); with blood-brain barrier (BBB) permeability, each molecule is given a digital score (1 = likely permeable; 0 = likely impermeable). A two-tailed Student’s T-test determined significance. The mean and the standard error of the mean are shown.

Second, we compared the *pgp-13*-inducers to the Spectrum molecules predicted to be PLD-inducing CADs. To identify PLD-inducing CADs, we employed a chemoinformatic formula that predicts whether a molecule is a CAD that will induce PLD in human cells based on the molecule’s lipophilicity and pKa (Pelletier *et al*. 2007) (see methods; Source Data file 1-tab 1, columns GL-GZ)). With this formula, we identified 288 Spectrum molecules that are predicted to be PLD-inducing CADs (Figure 3B; Source Data file 1-tab 1, column HA). This predictive formula identifies 73% of the gold standards (Figure 3B), which is 6.1-fold more than expected by random chance (*p*=3E-55). The predictive formula also identifies 73% of the *pgp-13* inducers (Figure 3B), which is also 6.1-fold more than expected by random chance (*p*=6E-169). Like the gold standards, the predicted PLD-inducing CADs that fail to induce *pgp-13* expression are significantly more polar and less likely to pass the skin barrier than those molecules that drive *pgp-13* expression (Figure 3C; Source Data file 1-tab 1). Importantly, the chemoinformatic predictor, which was trained on human data (Pelletier *et al*. 2007), does equally well with predicting which molecules will induce *pgp-13* expression in worms. This reinforces the idea that the properties of molecules that drive *pgp-13* expression in worms are intimately related to those that make a molecule a PLD-inducing CAD in humans.

### Most *pgp-13* Inducers Require NHR-70 and NHR-107

Above, we show that terconazole’s induction of *pgp-13* is dependent on both NHR-70 and NHR-107, while amiodarone’s induction of *pgp-13* is dependent on only NHR-70. We were curious to know whether most CADs require both nuclear receptors to induce *pgp-13* expression. We therefore assembled 195 molecules from our original 293 hits into a new mini-library (based on additional stringency and availability) (see bolded nodes of Figure 3A). We screened this mini-library for the ability of the molecules to induce *pgp-13*p::YFP expression in an NHR-70 and NHR-107-dependent manner. We found that 149 (76%) of the 195 molecules were dependent on both nuclear receptors to induce robust *pgp-13* expression (which we call ‘class I’ molecules) (*p*<0.05), 24 molecules (12%) were dependent on only NHR-70 (which we call ‘class II’ molecules) (*p*<0.05), and 3 (2%) were not dependent on either nuclear receptor (which we call ‘class 3’ molecules) (*p*<0.05) (Figure 4A; Source Data File 1-tab 1-columns ID-LP).

**Figure 4.**
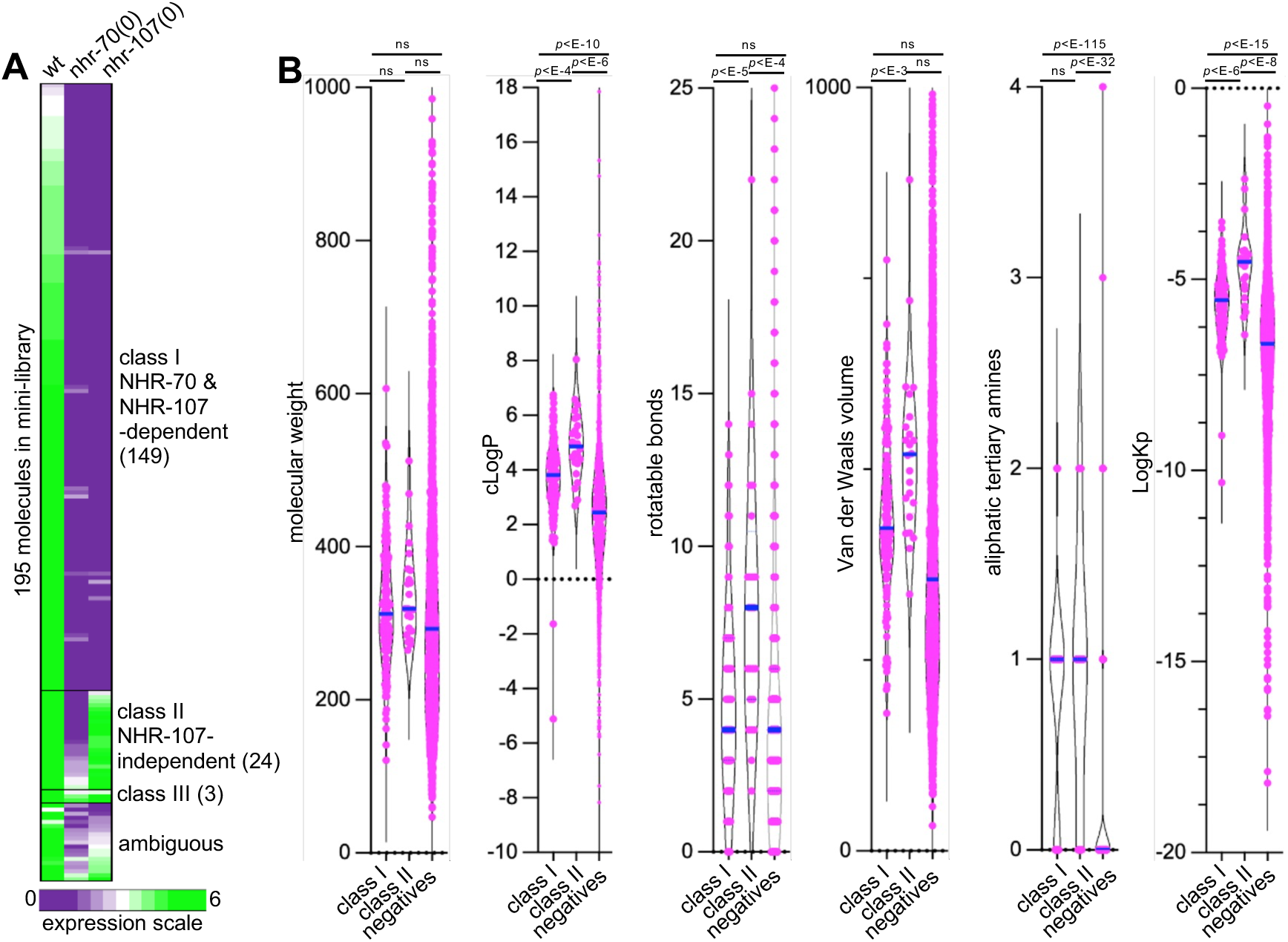
A Screen of 195 Small Molecules for their Ability to Induce *pgp-13* Expression in an NHR-70 and NHR-107-Dependent Manner. **A.** A heatmap showing the ability of 195 cherry-picked small molecules from the Spectrum library to induce *pgp-13p*::YFP expression in either a *wild type* (RP3507), *nhr-70(tm1697)* (RP3518) or *nhr-107(tm4512)* (RP3516) background at a concentration of 30 μM. 149 small molecules induced lower *pgp-13* reporter expression in the *nhr-70(0)* and *nhr-107(0)* mutants compared to control (*p*<0.05). 25 small molecules induced lower expression of the *pgp-13* reporter in only the *nhr-70(0)* mutant compared to control (*p*<0.05), and another 3 molecules did not have significantly lower expression in either mutant (*p*>0.05). See Source DataFile 1-tab 1- columns ID-LP for details. **B.** Violin plots of the indicated physicochemical properties for the class I and II molecules (indicated in A) relative to 2024 true negatives within the Spectrum library. The median is indicated. A two-tailed Student’s T-test determined significance. For clarity, 21, 1, 13, and 63 data points from the true negative class are left off from the molecular weight, cLogP, rotatable bonds, and van der Walls volume graphs, respectively (but are accounted for in the medians and statistical analyses).

Another 19 molecules did not fall into any of these bins based on statistical analyses and cutoffs. Physicochemical analyses show that class II molecules are enriched for molecules that are longer and less bulky than class I molecules; class II molecules are significantly (*p*<0.001) more hydrophobic (cLogP), more flexible (rotatable bonds), have a larger volume (van der Waals volume) without having a greater molecular weight compared to class I (Figure 4B; Source Data File 1-tab 12). Perhaps relatedly, class II molecules are more likely (*p*<0.001) to penetrate human skin than class I molecules (LogKp) (Figure 4B). These results provide a physicochemical basis for why different CADs may rely on a different combination of nuclear receptors to drive a CAD-response. Furthermore, there are a small number of CADs that drive *pgp-13* reporter expression independent of either nuclear receptor, suggesting that additional CAD-detection components beyond NHR-70 and NHR-107 may exist.

### Metabolic Analyses Reveals a CAD Detoxification System

The above results suggest that the CAD-induced expression of PGP-13 is part of a drug-response pathway and that there may be other genes like *pgp-13* whose expression is induced by the CADs in a nuclear receptor-dependent manner. To identify such genes, we examined published global expression analyses in response to the CADs mianserin and tamoxifen (Rangaraju *et al*. 2015; Diot *et al*. 2022). 21 genes, including *pgp-13*, were significantly and coincidentally up-regulated in both studies. We cross-referenced this gene list with a meta-analysis of 2243 genome-wide expression experiments that identified 11 genes that are significantly co-regulated with *pgp-13* (Schmauder AND Richter 2021). In total, five genes were common among the three datasets; *pgp-13*, three cytochrome P450s (aka P450s) including *cyp-35B1*, *cyp-35B2*, and *cyp-35B3*, and the predicted oxidoreductase *cror-1* (Figure 2C). CYP-35B1 is most similar to CYP-35B2 at the amino acid level (75%), while CYP-35B3 is least like the other two paralogs at 73% and 72% identity with CYP-35B1 and -35B2, respectively. The CYP-35B family members belong to P450 clan 2 and do not have a clear one-to-one orthology with homologues of human clan 2 P450s (Larigot *et al*. 2022), although human clan 2 member CYP2D6 is known to interact with positively charged molecules and can metabolize CADs (de Groot *et al*. 1999; Wang *et al*. 2015). We verified by qRT-PCR that these genes are indeed upregulated by multiple CADs in a manner dependent on either NHR-70 or NHR-70 and NHR-107 (Figure 2C; Supplemental Figure 5). We hypothesize that these five enzymes are part of a CAD defense system (Figure 2B). *cror-1* will not be further discussed here.

We investigated whether the components that are upregulated in response to CADs are involved in CAD detoxification. We first tested the hypothesis that NHR-70 is key in turning on machinery that metabolizes the CADs. To test this, we compared the ability of wild type worms to generate oxidized products of the CAD to that of the *nhr-70(0)* null mutant using liquid chromatography-coupled mass spectrometry (LCMS). For LCMS analyses, we focused on CADs that: i) induce robust expression from the *pgp-13p*::YFP reporter; ii) are structurally distinct; iii) have a tertiary amine and/or is predicted to be a PLD-inducing CAD; and iv) have a chloro group, which are not naturally found in animal metabolomes and exist as two highly abundant isotopes that are easily identified in LCMS analyses (Zhu *et al*. 2011). The chosen CADs are shown in Figure 5. Of the six CADs examined in detail, worms generated NHR-70-dependent oxidized metabolites above background levels for all molecules except terconazole (*p*≤0.05)(Figure 5a). This indicates that not all CADs that upregulate the CAD defense system are robustly metabolized by NHR-70-driven products.

**Figure 5.**
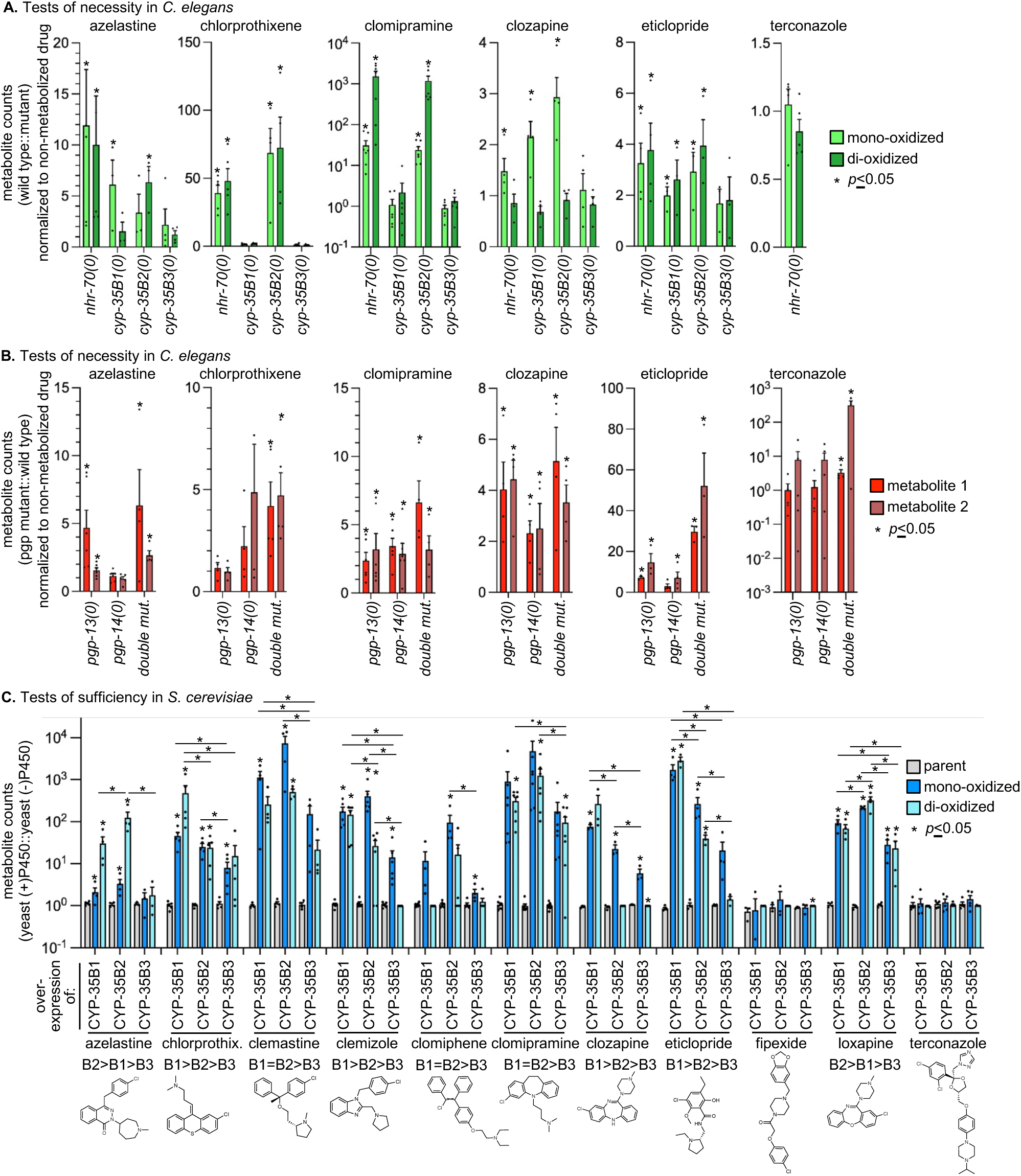
Metabolic Analyses of CADs. **A.** The abundance of mono or di oxidized products of the indicated parent molecule in the lysates of wild type worms relative to their abundance in the indicated mutants as measured by LCMS. The strains used are wild type (N2), TM1697 *nhr-70(tm1697),* RB2216 *cyp-35B1(ok2998)*, TM7435 *cyp-35B2(tm7435)*, and TM12255 *cyp-35B3(tm12255).* **B.** Similar to A, the abundance of metabolites that accumulate in the lysate of the indicated PGP mutant relative to the wild type. The strains used are RB894 *pgp-13(ok747)*, RP2960 *pgp-14(ok2660)*, and RP3495 *pgp-13(tr696) pgp-14(ok2660).* In both A and B, counts are normalized to the abundance of the parental molecule in each strain, and the exact masses of each metabolite and the *p* values (one tailed Student’s T-test relative to the normalized parent molecule counts) are found in Source Data File 1. N≥3 biological trials, n≥2 technical repeats. **C.** The abundance of the parent and mono and di oxidized metabolites in yeast (lysate and buffer combined) expressing the indicated P450 relative to (-) control yeast that do not express heterologous P450s. Chlorprothix., chlorprothixene. Statistical tests (one tailed Student’s T test) are done relative to parent mass, or when indicated with a horizontal line, relative to another P450-expressing strain. All *p*-values and exact masses are reported in Source Data File 1, tabs 14 and 15. In all graphs, the standard error is shown.

Next, we asked whether any of the three cytochrome P450s, whose expression is induced by CADs in an NHR-70-dependent manner, are necessary for CAD metabolism (oxidation). We again used LCMS to measure the accumulation of CAD metabolites in wild type worms compared to loss-of-function mutants of *cyp-35b1*, *35b2* and *35b3*.

We found that the loss of CYP-35B2 reduces the oxidation of all five CADs tested while the loss of CYP-35B1 reduces the oxidation of only three (*p*≤0.05) (Figure 5a). The loss of CYP-35B3 shows no decrease in oxidized CADs, consistent with the observation that the expression of CYP-35B3 is not induced by CADs as much as CYP-35B1 and B2 (Figure 2c). This data demonstrates that CYP-35B1 and -35B2 are necessary for robust CAD metabolism.

We next investigated the role of PGP-13 in CAD detoxification by asking whether CAD metabolites accumulate more in the *pgp-13(0)* null mutant relative to wild type.

Accumulation of CAD metabolites in the *pgp-13(0)* mutant would suggest a role for the PGP in drug efflux. Of the six CADs examined, we found multiple metabolites of four CADs that accumulate in the mutant (Figure 5b). Given that PGP-13 acts at least partially redundantly with PGP-14 in crystal formation (see above), we asked whether PGP-13’s role in CAD efflux for some drugs might be obscured by the activity of PGP-14. Indeed, the two CADs for which we failed to see CAD metabolite accumulation in the *pgp-13(0)* single mutant accumulate in the *pgp-13(0) pgp-14(0)* double mutant (Figure 5b). Hence, PGP-13 plays a role in CAD metabolite efflux both independently and redundantly with PGP-14, depending on the CAD in question.

We also investigated whether any of CYP-35B1, B2 or B3 are sufficient for CAD metabolism. To do so, we exploited our previously established methodology to heterologously express worm P450s in the yeast *S. cerevisiae* to determine their metabolic capability (Burns *et al*. 2023; Knox *et al*. 2024; Collins *et al*. 2025). Because the yeast experiments are less labour intensive compared to the worm metabolic analysis, we were able to examine more CADs. Of the 11 CADs examined, 9 were oxidized by at least one of the three P450s with CYP-35B2 metabolizing 9, CYP-35B1 metabolizing 8, and CYP-35B3 metabolizing 6 (*p*≤0.05) (Figure 5c). We were curious to determine whether the P450s inability to metabolize fipexide and terconazole in yeast might be related to their potential endogenous detoxification and/or a failure to robustly accumulate in yeast. We therefore compared the ratio of the accumulation of the CAD in the yeast cell lysate relative to what is found in the buffer for each of the 11 CADs. The only CAD that has a distinct accumulation pattern is clomiphene (*p*<0.05) (Supplemental Figure 6). Hence, the failure of fipexide and terconazole to be metabolized by yeast-expressed CYP-35B family members is not due to yeast detoxification.

CYP-35B1 is better able to metabolize chlorprothixene and eticlopride compared to CYP-35B2, which is seemingly contradictory to the worm mutational analysis shown in Figure 5a. However, *cyp-35b2* is expressed ∼32 fold more than *cyp-35b1* in worms (Figure 2c). In yeast, all 11 P450s are expressed using the same galactose-inducible promoter and are likely expressed at roughly equivalently levels. Hence, while 35B1 may be able to metabolize select CAD structures better than 35B2 in yeast, its lower expression in worms has less of an impact on CAD metabolism. Regardless, our data demonstrates that CYP-35B family members are both necessary and sufficient to metabolize CADs in *C. elegans*.

### Computational Modeling Reveals CYP-35B2 D311 as Key to in CAD Metabolism

Given the positive charge of CADs in cells, we were curious to know whether electrostatic interactions play a role in the ability of the CYP-35B family to metabolize CADs. Using CYP-35B2 and chlorprothixene and clomipramine as exemplar interactions, we employed structural modeling and molecular dynamics to model how charged and uncharged CADs interact with the P450. Protein structures, with the HEME and CAD, were predicted using Boltz-1 (Wohlwend *et al*. 2025) and then simulated with Amber 22 (Case *et al*. 2025). We found that the carboxylate of residue D311 is frequently within hydrogen bonding distance (<3 angstroms) of the charged moiety of the CADs, but is never within this distance when the uncharged molecules are modeled (Figures 6a and 6b; Supplemental Figure 7).

**Figure 6.**
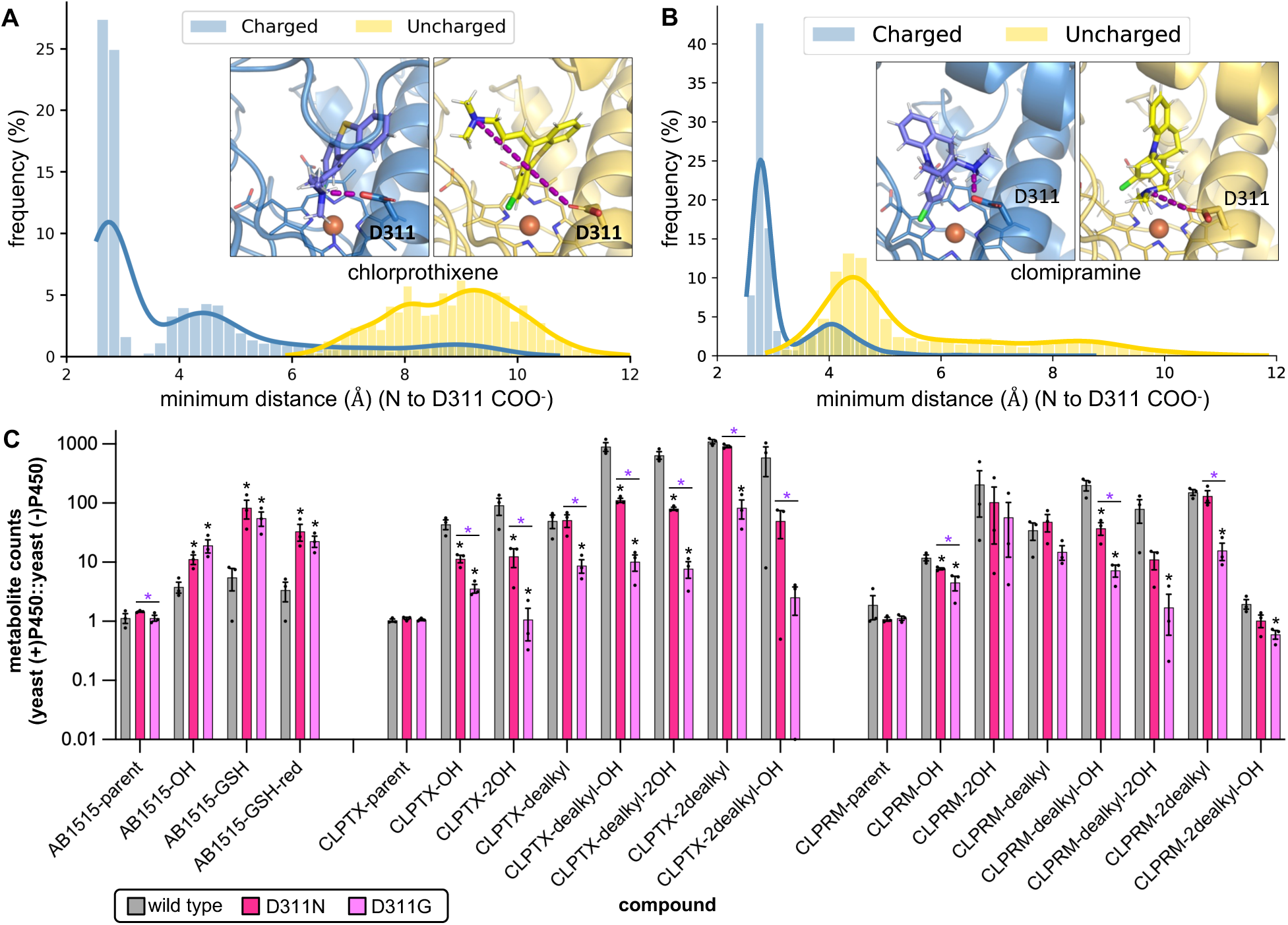
CYP-35B2 D311 is Important for CAD Metabolism. **A-B.** Computational modeling and simulation reveals that charged (blue) chlorprothixene (A) and clomipramine (B) frequently hydrogen bond with D311 while uncharged (yellow) versions do not consistently interact with D311. Representative conformations from molecular dynamics simulations are shown inset. **C.** The abundance of the indicated presumptive compound as identified by exact mass spectrometry as produced in yeast expressing either the wild type, D311N or D311G CYP-35B2. The abundance found in lysate and buffer was combined. Y axis is log_10_. N=3 biological trials, n=2 technical repeats. A one-tailed Student’s T test was used to measure differences. A black asterisk indicates *p*<0.05 when compared to the wild type; a purple asterisks indicates *p*<0.05 when comparing the two mutants. OH, hydroxylated metabolite; GSH, glutathione conjugate; GSH-red, reduced glutathione conjugate; dealkyl; dealkylated metabolite; AB1515, 5-(furan-2-yl)-3-(m-tolyl)-1,2,4-oxadiazole; CLPTX, chlorprothixene; CLPRM, clomipramine. The standard error of the mean is shown.

To test whether the anionic D311 residue of CYP-35B2 plays a role in cationic amphiphilic drug metabolism, we substituted D311 for the more conservative but uncharged asparagine residue, and for the less conservative glycine residue. We then used LCMS to ask what impact these substitutions have on the ability of CYP-35B2 to metabolize chlorprothixene and clomipramine in our heterologous yeast assay. We focused on exact masses consistent with the hydroxylation and/or dealkylation of the molecules. For most metabolites, both D311N and D311G produced less than wild type (*p*<0.05) (Figure 6c). Furthermore, D311G disrupts metabolism more than D311N for most metabolites (*p*<0.05) (Figure 6c), which is consistent with the divergence in structures of the two substitutions. To control for the possibility that the mutations simply make the enzyme less efficient, we compared the ability of the two mutant P450s to generate metabolites from a non-CAD tool compound that we discovered that we call AB1515 (5-(furan-2-yl)-3-(m-tolyl)-1,2,4-oxadiazole) relative to the wild type. The two mutants generated more AB1515 metabolites than the wild type (*p*<0.05)(Figure 6c). Hence D311 is likely key in mediating the interaction with CADs via electrostatic interactions.

### The CAD Detoxification Pathway Protects Against Pathogenesis

The results presented above suggests that a CAD-responsive pathway exists to protect worms from the potentially pathogenic effects of CADs. We tested this idea in two ways. First, we examined the viability of wild type worms grown in the presence of various CADs compared to relevant nuclear receptor and *pgp* mutants grown in the same compound using our standard 6-day viability assay (Burns *et al*. 2015). While *pgp-13* and *pgp-14* single mutants were hypersensitive to some CADs, the *pgp-13 pgp-14* double mutant showed more hypersensitivity to most CADs (Figure 7A). Consistent with our earlier results that show that PGP-13 expression can compensate for the loss of PGP-14 (Supplemental Figure 4; Figure 5B), these data suggest that PGP-13 and PGP-14 play redundant roles in the export CADs and their metabolites. Mutants of *nhr-70* and *nhr-107* are similarly hypersensitive to most CADs, but especially those where CAD metabolite production are particularly dependent on NHR-70, namely azelastine and clomipramine (Figure 7A).

**Figure 7.**
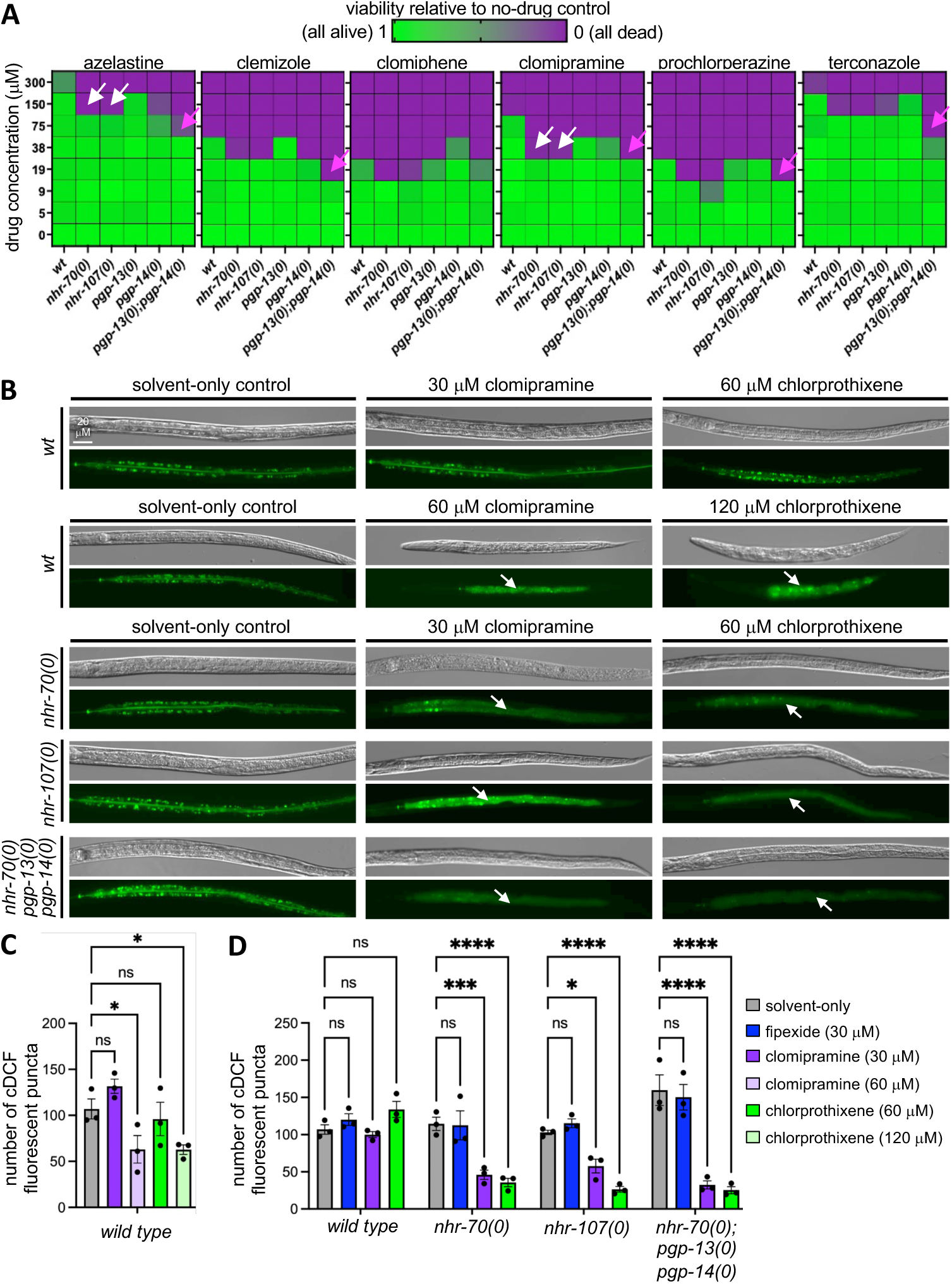
CADs Induce Pathology in *C. elegans*. **A.** Drugs assayed are indicated on the top, genotype on the bottom. Concentrations are the same for all drugs. Note the ∼4-fold hypersensitivity of azelastine and clomipramine in the nuclear receptor mutants (white arrows). Note the apparent redundancy of PGP-13 with PGP-14 in their presumptive role in drug efflux (pink arrows) (i.e., without the two pumps, the animals are more sensitive to the CADs, compared to either single mutant). N=3 biological trials and n=2 wells with 20 animals per well. **B.** Examples of cDCF signal in the indicated genotypes following 24-hour incubation with solvent or the indicated CAD. Note the diffuse fluorescence throughout the intestine (arrows) in the wild type animals treated with CADs at the higher concentrations or in the mutants treated with CADs at the lower concentrations. **C.** Quantification of puncta for the animals assayed in differing concentrations of the CADs. **D.** Quantification of puncta for the animals assayed in different mutant backgrounds. For both C and D, the legend is on the right, N=3 biological trials with n=10 worms/trial. Statistical significance was determined using a two-way ANOVA followed by Dunnett’s multiple comparisons test, where **** indicates *p*<0.0001 and * indicates *p*<0.05. Standard error of the mean is shown.

Second, we investigated whether *C. elegans* lysosome-related organelles are disrupted by CADs. We focused on the gut granules of the worm because they are well-characterized large acidic organelles that are terminal compartments for degradation and recycling endocytic pathways (Clokey AND Jacobson 1986; Bossinger AND Schierenberg 1996; Hermann *et al*. 2005). Previous work has employed carboxy-2’,7’-dichlorofluoresceine diacetate (cDCFDA) to study the pH of the gut granules (Baxi *et al*. 2017). cDCFDA is converted to the fluorescent cDCF (carboxy-2’,7’-dichlorofluoresceine) by esterases, which are robustly expressed in the gut (Edgar AND McGhee 1986). Because cDCF has a pKa of 4.2 with multiple ionizable groups, a significant fraction of cDCF remains deprotonated even in acidic compartments like the gut granules and consequently accumulates within the organelle because of its charge (Nedergaard *et al*. 1990).

CADs are well-known to neutralize acidic compartments and consequently disrupt their function (Nielsen *et al*. 2024). Because cDCF accumulation in acidic organelles is dependent on a pH difference within the organelle relative to the cytosol, we used cDCFDA to track gut granule neutralization. We performed a small dose-response analysis of two CADs (chlorprothixene and clomipramine) using the cDCFDA gut granule acidity marker. We found that 60 μM chlorprothixene and 30 μM clomipramine had an unaltered pattern of concentrated cDCF accumulation in gut granules like the solvent-only negative controls. However, when incubated in twice as much CAD, cDCF localization became less concentrated with the gut granules (Figure 7B-7C). These observations are consistent with the CAD’s ability to neutralize acidic organelles.

Given that NHR-70 and NHR-107 are key in the metabolism of chlorprothixene and clomipramine (Figure 5), we reasoned the corresponding loss-of-function mutants would increase the internal concentration of the parent CAD molecules and enhance the loss of cCDF signal within the gut granules. Indeed, we observed the same diffuse cDCF pattern in the gut with the 60 μM chlorprothixene and 30 μM clomipramine treatments in the *nhr-70(0)* and *nhr-107(0)* mutant backgrounds that were observed only at the higher concentrations in the *wildtype* background (Figure 7C-7D). These results further validate our model that the CAD-response machinery helps defend against CAD-related pathologies.

### Soil Microbes Elicit the CAD Defence Response

Above, we show that hundreds of diverse compounds trigger a robust CAD defense response in *C. elegans*, prompting the question of why the pathway exists at all. To address this, we asked whether bacteria in the natural environment of the worm might also trigger the expression of the CAD defense reporter. *Actinobacteria*, of which *Streptomyces* constitute the largest genus, are common within the natural environment of *C. elegans* (Kampfer 2006; Samuel *et al*. 2016), and produce an abundance of bioactive natural products that include molecules that we employ as antibiotics, anticancer therapeutics, immune suppressants, antifungals and other drugs (Ho AND Nodwell 2016; Daniel-Ivad *et al*. 2018). We screened 65 diverse *Actinobacteria* strains assembled from diverse locations around North America by scientists at Cubist Pharmaceuticals Inc (later sold to Merck). These strains produce diverse secondary metabolites, most of which are secreted into the media and/or deposited onto the surface of sporulating colonies (e.g. Supplemental Figure 8). From this collection, we identified 2 strains (1G9 and 1E6) capable of eliciting obvious *pgp-13p*::YFP expression (Figure 8). We tested whether this increase in signal was dependant on NHR-70 and NHR-107 and found that it was (*p*<0.005), indicating that the increase in *pgp-13* expression is likely elicited by a CAD-like natural product made by the bacteria (Figure 8). These observations suggest that a CAD defense pathway may have evolved in *C. elegans* to increases its survival rate in the presence of CAD-producing microbes.

**Figure 8.**
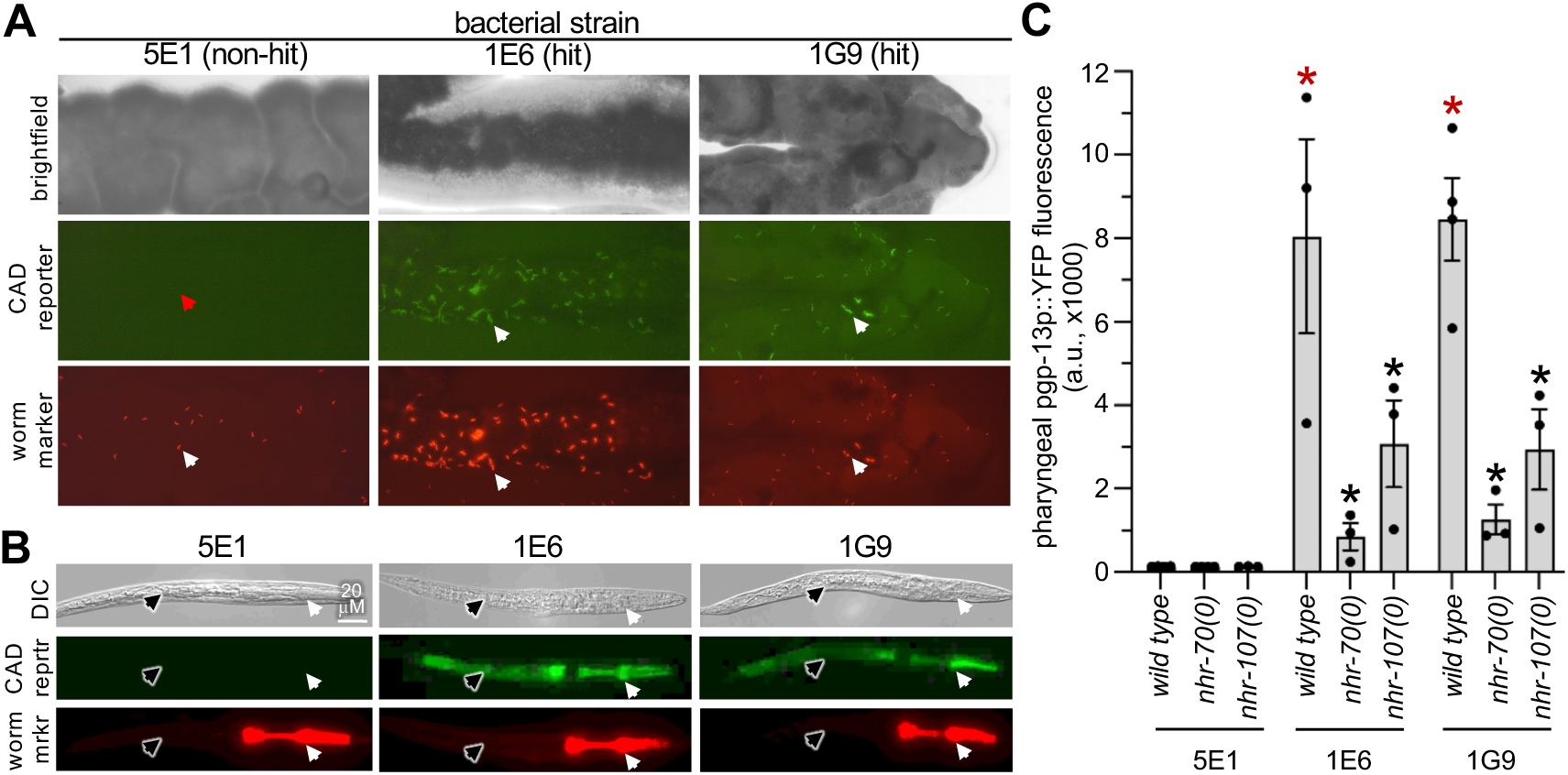
Induction of the CAD Defense System by *Streptomyces* Bacterial Strains. **A.** Examples of one non-hit (5E1) and two hits (1E6 and 1G9) from our screen for *Streptomyces* strains that can elicit YFP expression from the RP3507 *trIs114* [*pgp-13p::YFP; p382(myo-2p::mCherry*)] CAD reporter strain after 24 hours of co-incubation. The top panels are brightfield images of the bacterial colony, the middle panels show the YFP signal (white arrows show one example) or lack thereof (red arrow), and the bottom panels show the constitutive expression of mCherry in the pharynx that marks the presence of the worms. **B.** Representative high magnification images of individual worms exposed to the three bacterial strains. The white arrowheads indicate the pharynx; the black arrowheads indicate the intestine. **C.** Quantification of the YFP signal in the indicated genetic background grown in the indicated *Streptomyces* strain for 24 hours. N=3 biological trials with n=10 worms/trial. A two way ANOVA was used to determine statistical differences. Red asterisks indicate *p*<0.0001 relative the corresponding genotype grown on 5E1; black asterisks indicate *p*<0.005 relative to wild type within the same growth conditions.

## Discussion

Here, we demonstrated that *C. elegans* harbours a pathway that defends against cationic amphiphilic small molecules (aka CADs). The pathway is adept at responding to a wide variety of CAD structures. CAD exposure upregulates the expression of CAD-metabolizing cytochrome P450s and PGP-13, which acts partially redundantly with its paralog PGP-14 to export CAD metabolites. Without an intact CAD defense pathway, the animal is less able to protect itself against the effects of CAD accumulation. Given the unclear orthology between components of this pathway and human homologs, how these insights will translate to CAD pharmacokinetics in humans remains to be determined. However, the findings indicate that animals can indeed mount a robust and coordinated defense against CADs that have phospholipidosis-inducing potential.

Evidently, PGP-13 acts as a multidrug CAD metabolite efflux pump but can also weakly compensate for the loss of PGP-14, which likely exports phospholipids (Kamal *et al*. 2023). The relationship between PGP-13 and PGP-14 is remarkably similar to that of human MDR1 (aka ABCB1) and ABCB4; the MDR1 multidrug export pump is also capable of exporting phospholipids but less effectively than ABCB4 (van Helvoort *et al*. 1996) (Raggers *et al*. 2001). Furthermore, PGP-13 and PGP-14 directly neighbor one another in the genome, as do MDR1 and ABCB4 (Supplemental Figure 2). Whether the conservation of the physical relationship between the paralogous pairs is coincidental or functional remains to be determined.

One goal of this study was to determine whether small molecules could be used to suppress the phenotypes associated with the loss-of-function of *pgp-14*, which has the potential to serve as a model for ABCB4 deficiency in humans. The only drugs recovered from our screen for *pgp-14(0)* suppressors were the CADs, which as described above, can rescue the loss of PGP-14 by upregulating the expression of its paralog PGP-13.

Whether CAD treatment in humans might induce the expression of the ABCB4 paralog ABCB1 or some other phospholipid efflux pump and provide some measure of relief for PFIC3 patients with ABCB4 hypomorphic mutations is an open question. That ABCB1 is drug-inducible in hepatocytes (Williamson *et al*. 2013; Jarvinen *et al*. 2023) provides some encouragement for the hypothesis, but the complications surrounding the use of phospholipidosis-inducing drugs is a potential confounding issue. Investigating whether CADs can suppress the phenotypes of ABCB4 deficiency in *mdr2* mutant mice (Weber *et al*. 2019) would be one step towards testing this hypothesis.

Many more Spectrum library small molecules induce PGP-13 expression than rescue the *pgp-14(0)* null mutant phenotype. Three reasons may explain this discrepancy. First, the temporal kinetics of CAD rescue of *pgp-14(0)* is slow. We observe only small crystals after multiple days of coincubation of the mutant with the CADs plus a crystalizing molecule. By contrast, we see crystals within minutes of coincubation of a crystalizing molecule with wild type. Hence, a CAD that may be a slightly weaker PGP-13 inducer might be missed in the *pgp-14(0)* suppressor screen. Second, some CADs that induce PGP-13 expression are toxic and confound *pgp-14(0)* suppression. Finally, many CADs that we identified as *pgp-13* inducers elicit expression more robustly in the intestine as opposed to the pharynx, which will reduce their ability to suppress the *pgp-14(0)* phenotype, which is restricted to the pharynx.

It remains an open question as to whether CADs interact directly with the NHR-70 and NHR-107 nuclear receptors to induce a transcriptional response. Given the extensive diversity of CADs that induce the pathway, this model would require that the ligand binding domains of NHR-70 and NHR-107 be equivalently plastic in their interaction with ligands as the P450s that they regulate. Alternatively, the nuclear receptors may be responding to an endogenous metabolite generated as a secondary consequence of CAD accumulation. Using our LCMS protocol that we optimized for small molecule detection, we did not observe a mass that reproducibly accumulates in worms treated with different CADs relative to controls. However, it remains possible and perhaps even likely that endogenous metabolites that accumulate in CAD-treated worms are not captured by our current methodology. Future studies may shed light on this issue.

We demonstrated that the expression of CYP-35B family members 35B1, 35B2 and 35B3 is induced by CADs. Ethanol, caffeine, ethidium bromide, PCB52, polycyclic aromatic hydrocarbons and several pesticides, none of which are CADs, are also known to induce CYP-35B expression (Menzel *et al*. 2001; Larigot *et al*. 2022). Five of these molecules (caffeine, chlorpyrifos, dichlorvos, rotenone, and thiabendazole) are included in the Spectrum small molecule library and were all negative for the induction of *pgp-13* expression in our screen. Hence, the CYP-35B family members must have multiple distinct regulatory inputs in response to different xenobiotics.

We have also demonstrated that CYP-35B1 and 35B2 are necessary for CAD metabolism in worms and that all three P450s are sufficient for this activity upon being expressed in yeast. Metabolic studies of *C. elegans* P450s are sparse (Larigot *et al*. 2022) and we could not find evidence within the literature of previously characterized metabolic reactions involving CYP-35B family members. However, our finding that our tool molecule AB1515 is metabolized by CYP-35B2 indicates that at least CYP-35B2’s substrate selectivity extends beyond CADs. Our results also suggest that CYP-35B2’s D311 contributes to a filtering mechanism likely by increasing the affinity to CADs via electrostatic interactions whilst at the same time reducing interactions to non-CAD molecules like AB1515.

The only CADs that the CYP-35B family failed to metabolize are fipexide and terconazole, despite these molecules being able to robustly induce the marker of CAD defense (see Figure 4a and Source Data file 1-tab 1). There may be multiple reasons why these CADs are not metabolized. First, azoles such as terconazole are known to inhibit P450s (Aoyama *et al*. 1984; Yoshida AND Aoyama 1984), making it possible that terconazole disrupts CYP35B function. Second, fipexide has the lowest pKa (6.09) of the 11 molecules tested and barely makes the CAD cutoff score (see Source Data file 1-tab 1). Hence, fipexide is likely a weaker interactor with the CYP-35B family members given that the positive charge of CADs is an important determinant of their metabolism by these P450s. Finally, the shape of fipexide and terconazole sets them apart from the remaining CADs tested in yeast; they harbour the longest atom chains among the 11 molecules. Regardless, it is clear that there is not a one-to-one correspondence in being able to engage the CAD detection machinery and being subject to its defenses.

Obviously, the CAD detoxification pathway in *C. elegans* did not evolve to defend against anthropogenic CAD molecules that have existed for fewer than 100 years. Indeed, our small-scale screen of 65 *Streptomyces* revealed multiple strains that induced the CAD defense reporter, suggesting that natural products engage the CAD defense pathway. Furthermore, we know that individual *Streptomyces* strains generate different types of secondary metabolites depending on their growth conditions (Yoon AND Nodwell 2014; Saito AND Arai 2024), suggesting that we have only skimmed the surface in identifying strains capable of engaging the *C. elegans* CAD defense pathway. At the very least, our observations provide a framework for rationalizing the existence of a pathway that was discovered using anthropocentric-derived molecules.

## Methods

### *C. elegans* husbandry, screening and microscopy

Unless otherwise noted, the following canonical alleles were used in genetic analyses: The N2 *wild type* Bristol strain, RB2216 *cyp-35B1(ok2998)*, TM7435 *cyp-35B2(tm7435),* TM12255 *cyp-35B3(tm12255),* TM1697 *nhr-70(tm1697),* TM4512 *nhr-107(tm4512),* RB894 *pgp-13(ok747)*, RP2960 *pgp-14(ok2660)*, and the RP3495 *pgp-13(tr696) pgp-14(ok2660),* RB894 *pgp-13(ok747)*, RP2960 *pgp-14(ok2660),* RP3522 *nhr-70(tm1697)*; *pgp-14(ok2660), and* RP3523 *nhr-107(tm4512)*; *pgp-14(ok2660)* double mutants. All strains were cultured using standard techniques (Lewis AND Fleming 1995) at 20°C. The six-day liquid viability assays were performed as described previously (Burns *et al*. 2015). Briefly, *E. coli* (HB101) was cultured in Lysogeny broth (LB) media at 37°C overnight to make saturated culture. The cells were collected by centrifugation (3,200 rpm for 20 minutes) and 2-fold concentrated by resuspending them in 0.5 volume of nematode growth medium (NGM; 3 mg/mL NaCl, 2.5 mg/ml peptone, 5 mg/mL cholesterol, 1 mM CaCl_2_, 1 mM MgSO_4_ and 25 mM KH_2_PO_4_). 80 μL of HB101/NGM mix was dispensed into each well of a 96-well plate. Then compounds (typically 300 nL of 10 mM stock) or solvent-only (DMSO) control are added to the 96-well plate using a 96 pinner pinning tool according to the manufacturer’s instruction (V&P Scientific).

Approximately 20 first larval stage (L1) animals in 20 μL of M9 buffer (6mg/mL Na_2_HPO_4_, 3mg/mL KH_2_PO_4_, 5mg/ml NaCl, 2mM MgSO_4_) were then dispensed to each well of the 96-well plate. 96-well plates are incubated in a 20°C shaker at 200 rpm for six days. Unless otherwise indicated, all samples are assayed in technical duplicates for every run.

### The Screen and Initial Characterization of *pgp-14(0)* Small Molecule Suppressors

The objective of this screen was to identify small molecule suppressors of the *pgp-14(0)* mutant phenotype. Normally, wild type animals succumb to the lethal effects of crystal formation of the wact-190 small molecule within the pharynx cuticle; *pgp-14(0)* mutants resists these effects (Kamal *et al*. 2019; Kamal *et al*. 2023). Hence, we endeavored To identify small molecules that resensitize *pgp-14(0)* mutants to the lethality and crystal formation conferred by wact-190, we grew N2 wild type animals and *pgp-14* mutants *tr458* and *tr477* (both loss-of-function mutations (Kamal *et al*. 2023)) in liquid format in 96-well plates in duplicate (see the description of the 6-day viability assay above). With a subset of the library, mutants with the *pgp-14(ok2660)* deletion allele were also screened. N2 was treated with 30 µM of the screening compounds from Microsource’s Spectrum Library (Source Data File 1-tab 1) while the *pgp-14* mutants were treated with 30 µM of the screening compounds and 15 µM of wact-190 (Chembridge Inc). Our 2015 Spectrum Library contains 2560 clinically approved drugs and natural products (Source Data File 1-tab 1). For the screen, the number of adults (A) and larvae (L) was manually counted in each well for up to 50 animals. Beyond that, wells were given a score of ‘overgrown’ or OG (see OG, A and L scores in Source Data File 1). A series of retests, where a ceiling of 50 animals were counted, revealed 14 Spectrum molecules that induced lethality in the *pgp-14 null* mutant at 30 µM when co-incubated with 15 µM or 30 µM of wact-190 (see column ‘Z’ in Source Data File 1-tab 1). None of the Spectrum hits induced obvious phenotype in the absence of wact-190. We also tested whether candidate suppressors could restore wac-190 crystal formation in the *pgp-14(0)* (but not induce crystal formation in *wild type* animals) using previously-described methodology (Kamal *et al*. 2019). Briefly, 50 synchronized L1 animals were incubated in 30 µM of the Spectrum small molecule hits and either 30 µM of wact-190 or solvent-only control (1% DMSO) for 48 hours. Then, animals were transferred to a microcentrifuge tube and spun down at 2,800 rpm. After aspiration, 2 μL of 50 mM levamisole was added to 10 μL worm pellet. The worm pellets were placed on 3% agarose pad on glass slides and cover-slips were applied. A Leica DMRA 2 compound microscope was used at 40X magnification to investigate the presence of crystals in the anterior pharynxes of the worms. Openlab software was used to take DIC and birefringence images.

### A Forward Genetic Screen for Mutants that Suppress the Ability of CADs to Restore *pgp-14(0)* Sensitivity to the Lethal Effects of wact-190 Crystal Formation

The forward genetic screen mutagenesis was conducted as described in previous work (Burns *et al*. 2006). Briefly, RP3412 *pgp-14(ok2660); trIs104[p1138(pgp-14p::SMS-5B(genomic)::FLAG::mCherry); p1131(pgp-14p::YFP)*] parental (P0) worms were mutagenized in ethyl methanesulfonate at a concentration of 50 mM for 4 hours. We included extra copies of functional SMS-5 in the genetic background (i.e. *trIs104*) because we knew that mutations in *sms-5* would suppress the ability of the CADs to resensitize *pgp-14(0)* to wact-190 crystal formation because SMS-5 is necessary for crystal formation (Kamal *et al*. 2019). 100 synchronized and mutagenized F2 (second-generation) L1 worms were dispensed into each well of a 96-well plate, with a total volume of 50 µL containing 30 µM terconazole (CAS# 67915-31-5; Sigma-Aldrich) and 30 µM wact-190 suspended in liquid nematode growth media (NGM) with HB101 *E. coli* cell as a food source. The plates incubated over 14 days at 20°C in a shaking (200 rpm) incubator. Plates were then examined for any wells containing adults and progeny, which was indicative of the presence of a mutant that resists the activity of the CAD’s (terconazole) (i.e., resists the CAD’s ability to resensitize the *pgp-14(0)* mutant to the lethal effects of wact-190). These mutants were then isolated (ultimately only one line per resistant well) and cloned to enable further characterization.

To identify the mutant genes that were likely responsible for CAD-resistance, genomic DNA was prepared, and sequences were analyzed as previously described (Kwok *et al*. 2006). In brief, 100 µL of worms were collected in a 15 mL conical tube and washed three times with M9 buffer, followed by a single wash using phosphate-buffered saline (PBS). Excess media was aspirated without perturbing the worms, followed by flash freezing via liquid nitrogen and processing using the QIAGEN DNeasy Blood and Tissue Kit (Cat # 69504). DNA library construction and sequencing was conducted via the services of the Centre for Applied Genomics (Toronto) services. In brief, the 150-bp paired-end sequencing reads were examined for quality via FastQC (Andrews 2010), and Trim Galore! was used to cut off low-quality reads as well as adapter sequences. The processed reads were then aligned via the BWA-mem tool to the *C. elegans* N2 reference genome – release WS282 (Li AND Durbin 2009). Sequence alignments were sorted by coordinate order, and duplicate reads were marked and removed using Picard tools (https://broadinstitute.github.io/picard/). Variants were called and consolidated utilizing GATK’s HaplotypeCaller and CombineGVCFs (McKenna *et al*. 2010). Variants were then filtered on the following criteria to ensure high-confidence variant calls: Fisher Strand (FS) > 60.0, Mapping Quality (MQ) < 40.0, Mapping Quality Rank Sum (MQRankSum) < -12.5, Qual By Depth (QD) < 2.0, Quality Score (QUAL) < 30.0, Read Position Rank Sum (ReadPosRankSum) < -20.0 or < -8.0, and Symmetric Odds Ratio (SOR) > 3.0. Variants were subsequently annotated using Annovar to generate a list of exonic variants, from which candidate genes were identified (Wang *et al*. 2010).

### RNA Extraction, cDNA Synthesis and Quantitative Real-Time PCR

Synchronized L1 worms were treated with either vehicle (1% DMSO) or small molecule in 1% DMSO for 96 hours in liquid NGM with HB101 *E. coli* as a food source at a final concentration of two worms per microlitre in a total of 8 mL. RNA extraction was performed as described in (Riccio 2019). RNA within the aqueous portion of TRIzol™ (Invitrogen) was purified utilizing an Aurum™ Total RNA Mini Kit (Bio-Rad), and treated with TURBO™ DNase (Thermo Fisher) to remove genomic DNA. cDNA synthesis was conducted from purified RNA using the Applied Biosystems™ High-Capacity cDNA Reverse Transcription Kit (cat#4368814). Primers for targets were designed to span at least 1 exon-exon junction using NCBI Primer Blast (Ye *et al*. 2012). SsoAdvanced™ Universal SYBR Green Supermix (Bio-Rad) was utilized with 5 ng of cDNA and primers at 500 nM of each primer to conduct q-RTPCR on the QuantStudio™ 6 Flex Real-Time PCR System. The data was analyzed utilizing the comparative C_T_ method normalized to the common standard *rpl-33* (Schmittgen AND Livak 2008; Tao *et al*. 2020).

RPL-33: F 5’-GGCCCACAACAAGACTTTGAAAACC-‘3; R 5’-ATGGGTAGAGAAGCACGCGG-3’

PGP-14: F 5’-CACGACCAAATTGAACGACAGTATG-3’; R 5’-TGTAGTTGAAGTCATTTGTCTGGC-3’

PGP-13: F 5’-TTTCGCTACTGCAATTGTTCCGC-3’; R 5’-ACACATTTGTCATTCGACCCGAG-3’

CYP-35B1: F: 5’-GCTGAACACGAGATGTGCCG-3’; R: 5’-ACGTTTTCCGACGAGCAGAGA– 3’

CYP-35B2: 2. F: 5’-GGGCCAGTATCTCTTCCACTT-3’; R: 5’-GTGAAAACATCTCCATAAATCTGTC-3’

CYP-35B3: F: 5’-GGATGAGCTGAATGCAAGATGTGC-3’; R: 5’-CACTGCCAACAGTTAGGTCAAAG-3’

CROR-1: F: 5’-GTGGGCAACATGCGCAAC-3’; R: 5’-TTGTAGATGTTTTCCACCGACTC-3’

CROR-2: F: 5’-GGGCGTTTGAGGAGATGATG-3’; R: 5’-CTTACGATCAGCCAAGAACGC-3’

### Construction and analyses of *pgp-13* transcriptional and translational reporters

The pPRLT1187 *pgp-13p*::YFP transcriptional reporter was constructed by PCR amplifying 1.5 kb of the *pgp-13* sequence 5’ to the ATG using the 5’ primer 5’-aagtcTCTAGAattatgaaagaaatccgagaggtctg-3’ and the 3’ primer 5’-gtactgACCGGTcagcggttcgttttatacaattgtg-3’ from wild type (N2) genomic DNA, and then cloning the PCR product into a modified construct from the Andy Fire vector collection kit (L4666). Animals were made transgenic for this construct through microinjection into wild type (N2) animals and then chromosomally integrated using the ultraviolet radiation technique (Mello AND Fire 1995), resulting in *trIs114* [*pgp-13p::YFP; p382(myo-2p::mCherry*)].

The mNeonGreen::PGP-13 N-terminal fusion translational reporter was built with the help of *In vivo Biosystems Inc* by using CRISPR to insert mNeonGreen coding sequence, containing three synthetic introns, in-frame with a 30 bp linker (GGTTCGGGCTCGGGTTCTGGAAGTGGTAGT) immediately 5’ to the initiator methionine codon of the *pgp-13* gene, resulting in the integrated transgene *trIs115*. The sgRNA guides included the 5’ sgRNA GATGCCACCACAAGAAAACA and the 3’ sgRNA TTTTCTTGTGGTGGCATCAG. Importantly, the *trIs115* mNeonGreen::PGP-13 fusion gene was created in the background of the *pgp-14(ok2660)* deletion allele. We used this background to determine whether the resulting fusion protein was functional; the only known phenotype at the time of reporter construction was that *pgp-13*’s loss-of-function is capable of suppressing the ability of CADs to resensitize *pgp-14(0)*’s resistance to the lethal effects of wact-190 crystals. Furthermore, *pgp-13* and *pgp-14* are neighboring genes and recovering a *pgp-13* knock-in recombinant with a neighboring *pgp-14(0)* mutant would be highly unlikely. Hence the need for the creation of the mNeonGreen knock-in into *pgp-13* in the background of a *pgp-14(0)* mutant. The key result that demonstrated that mNeonGreen::PGP-13 is functional is that the RP3561 strain (which encodes the mNeonGreen::PGP-13 fusion in the background of the *pgp-14(ok2660)* null) yields crystals in the background of the CAD terconazole (which suppresses *pgp-14(0)* because it upregulates *pgp-13* expression). This outcome is identical to that of wild type *pgp-13* in the background of *pgp-14(0)*. If *pgp-13* had lost function through the fusion of mNeonGreen, terconazole would not have been able to resensitize *pgp-14(0)* to wact-190 crystals. See Supplemental Figure 4 for details.

### Analyses of *pgp-13* temporal expression and dose-response

The temporal and dose-response analyses of *pgp-13p*::YFP reporter expression was done in 96-well plate format with either a wild type background (strain RP3507) and/or the backgrounds of the *nhr-70(tm1697)* and *nhr-107(tm4512)* null mutants (strains RP3518 and RP3516 respectively). For each experiment, 3 independent biological repeats were performed with ∼150 worms per well. For the dose-response analyses, at least 12 technical repeats for each data point were prepared for each trial. For the temporal analysis at least 36 technical repeats were prepared for each trial. For each, ∼150 synchronized L1 worms were added to each well. Compound was added to the wells using a 96 pin pinner to a final concentration of 0, 15 or 30 μM, with a final DMSO concentration of 1%. Plates were incubated at 20°C for 48 hours on a shaking incubator. YFP and RFP signal was measured with a CLARIOstar fluorimeter (BMG Labtech Inc). Only YFP signal was considered in the analyses because fluctuations in basal RFP expression threw YFP::RFP ratio wildly, while YFP-only signal yielded reproducible signal patterns.

### Analyses of *pgp-13* Localization

Characterization of the expression pattern of the mNeonGreen::PGP-13 fusion reporter (RP3561 *trIs115* [mNeonGreen::PGP-13; *pgp-14(ok2660)*]) in the head of the animal was conducted by incubating ∼100 synchronized L1 worms in either solvent control or 30 µM of terconazole for 48 hours in liquid nematode growth media (NGM) with HB101 *E. coli* cells (OD600 ≥ 1.2) as a food source. Following the incubation on a tube rotator, worms were given the chitin stain calcofluor white at a final concentration of 104 µM for 1 hour and then imaged. Intestinal expression was investigated by incubating ∼100 synchronized L1 RP3561 animals in liquid nematode growth media (NGM) with HB101 *E. coli* cell as a food source with solvent or 30 µM of terconazole for 72 hours. Following the incubation, worms were incubated in 1mg/mL Texas Red-dextran (40,000 MW) for 1.5 hours. Imaging was performed using a Leica DM6B microscope equipped with 63x and 20x objectives.

### A Screen of the Spectrum Library for Small Molecules that Induce *pgp-13* Expression

*The Spectrum Screen for Small Molecule Inducers of pgp-13 Expression:* To identify small molecule inducers of *pgp-13* expression, we screened the RP3507 *trIs114* [*pgp-13p::YFP; p382(myo-2p::mCherry*)] animals against 2400 molecules of the Spectrum Library (Microsource Inc) (Source Data File 1-tab 1). We deposited synchronized ∼150 L1s into the wells of a 96-well plate along with 30 μM (final concentration) of compound in 1% DMSO and *E. coli* (an OD600 of ∼1.5) as a food source and incubated at 20 °C for 48 hours on a shaking incubator. Duplicate plates were set up and analyzed independently. The fluorescence signal from the pharynx and intestines of the worms were scored independently and manually binned into one of seven categories: 6, saturated; 5, very bright; 4, bright; 3 medium bright; 2 obvious signal, but not bright; 1 signal just above background; 0 no obvious signal. Given that the correlation between the two manual counts was +0.96 (Source Data File 1), we were confident in this scoring system. For the purposes of this analysis, we considered any molecule that resulted in an average score in the pharynx and/or the intestine of 2.5 or more to be a consistent inducer of *pgp-13* reporter expression.

### Generating a Structural Similarity Map

Molecule scaffolds were generated from Chemical SMILES of the Spectrum library were provided by the vendor (Microsource Inc). SMILES were edited to remove salts, and then reduced to their core scaffolds using the Strip-it command line tool version 1.0.2 (RINGS_WITH_LINKERS_1 scaffold definition). The resulting scaffold molecules were analyzed for pairwise structural similarity by calculating the Tanimoto coefficient of shared FP2 fingerprints using OpenBabel (http://openbabel.org) and a structural similarlity score of 1 (identical) was used to generate the structural similarity map, which was visualized using Cytoscape version 3.10.3 (Shannon *et al*. 2003).

Physicochemical properties of molecules were downloaded from SwissADME (Daina *et al*. 2017) and investigated for differences among different groups of molecules.

### Establishing a Predictive Model of Phospholipidosis(PLD)-inducing CADs

We constructed several predictive models of PLD-inducing CAD molecules based on the published model (Pelletier *et al*. 2007), each using a different method that predicts hydrophobicity of a small molecule (i.e., cLogP). We estimated the pKa of the most basic group of each molecule by using the MolGpKa API (Pan *et al*. 2021). The original MolGpKa API python script was modified to allow iteration through lists of smiles, and Microsoft Excel (v 16.43) was then used to deconvolute the output. We evaluated the different models against a suit of 112 gold standard CAD-inducing molecules identified from the literature (see Source Data File 1-tab 1), and then evaluated each based on their capture rate of the gold standards using the hypergeometric *p* value calculation.

The ‘PLD-inducing-CAD’ predictive model that captured the highest fraction of gold standards as a factor of the number of predicted PLD-inducing CADs (6.1 fold more than expected by chance alone; hypergeometric *p*-value E-55) was produced using the MLogP model (SwissADME) (Source Data File 1-tab 1). We therefore proceeded with our analyses using this model.

### Spectrum MiniLibrary Analyses

From the 293 Spectrum Library hits that induce *pgp-13p*::YFP expression, we selected those that induced robust expression in the pharynx (an average score of 3 or more) and was plentiful within our cherry-picking stock plates), leaving a total of 195 hits to analyze in more detail (bolded nodes in Figure 3a). These 195 molecules were re-arrayed into new 96 well plates and screened with the strains RP3507 *pgp-13p::YFP*, RP3516 *pgp-13p::YFP; nhr-107(0)*, and RP3518 *pgp-13p::YFP; nhr-70(0)* as described above with synchronized L1s grown for 48 hours at 20°C in a shaking incubator. The wild type (RP3507) strain was screened two independent times (N=2) with two technical repeats each (n=2); the two mutant strains were each screened four independent times (N=4) with two technical repeats each (n=2). Although we collected are reported data for expression in both the pharynx and intestine, we focused on only the pharynx because low background signal.

### Sample Preparation of *C. elegans* for Mass Spectrometry Analyses

30,000 synchronized L1 *C. elegans* were placed into 1.5 mL microcentrifuge tubes in 495 µL M9 buffer per sample. Compounds were added to achieve the desired final concentration, where DMSO final concentration is at 1% (v/v). Samples were incubated on a tube rotator to ensure proper at room temperature for 24 hours. Following the incubation, worms were transferred to 96-well filter plates (Pall P/N 8684) and the buffer was removed via vacuum filtration. Worms were then washed with 600 µL of M9 buffer and then resuspended in 50 µL of water and immediately frozen at -80 °C.

Frozen worm pellets were thawed and resuspended in 400 µL of butanol (BuOH) with the addition of 100 µL of 0.5 mm glass beads. Samples were lysed using a bead-beating device (Bead Ruptor 24, Omni International, Kennesaw, GA, USA), followed by centrifugation at 14,000 rpm for 10 minutes. 350 µL of the BuOH extract was then transferred to a new Eppendorf Safe-lock tube (cat. #022363204), and 350 µL of methanol (MeOH) is added to the original vial which was then bead-beaten a second time. After centrifugation at 14,000 rpm for 10 minutes, 350 µL of MeOH extract was combined with the BuOH extract in the new tube. The combined extracts were then dried at 60°C utilizing an Eppendorf Vacufuge (V-AL a pre-set mode for drying alcohol-based solutions). Dried samples were then stored at -20 °C until further processing.

The dried extracts of worm lysate and buffer (and yeast lysate and buffer-see below) were resuspended using 100 µL of a 50:50 mixture of LCMS grade acetonitrile and water. Samples were then vortexed for at least 30 seconds, followed by sonication for 15 minutes and centrifugation at 14,000 rpm for 15 minutes. For LCMS analysis, 25 µL of the supernatant was transferred into an amber vial insert and the remainder was stored at -20 °C.

### Expression of Wild Type and Mutant *C. elegans* CYP-35B P450s in the Yeast *S. cerevisiae*

The *C. elegans cyp-35b1*, *cyp-35b2*, and *cyp-35b3* cDNA sequences were obtained from WormBase (version WS296) and were codon-optimized for yeast expression. The codon-optimized ORFs were synthesized by Twist Biosciences and placed into the SpeI and HindIII restriction sites of the ATCC p416 GAL1 expression vector (URA3 selection marker; CEN6/ARSH4 origin of replication) (Montalibet AND Kennedy 2004). The *S. cerevisiae* stain RP4400 (S228c background) MATa his3Δ leu2Δ ura3Δ lys2Δ pdr5Δ snq2Δ yor1Δ can1::pGAL1-Ce emb-8 was transformed using the lithium acetate/single stranded carrier DNA/PEG method (Gietz AND Schiestl 2007) with the *cyp-35B1*, *cyp-35B2*, *cyp-35B3*, and empty vector negative control plasmids and were selected for on SD (-)URA medium. For the mutants, codon-optimized DNA fragments encoding *cyp-35B2* variants carrying the D311N or D311G substitutions were synthesized by Twist Bioscience with 5’ and 3’ adaptors: 5’-CAAAAAATTGTTAATATACCTCTATACTTTAACGTCAAGGAGAAAAAACCCCGGATTCTAGAACT AGT-3’, and 5’-AAGCTTATCGATACCGTCGACCTCGAGTCATGTAATTAGTTATGTCACGCTTACATTCACGCCCTCCCCCCACATC-3’, respectively, whose sequence corresponds to the multi-cloning site of the aforementioned ATCC p416 GAL1 expression vector. The fragments were co-transformed along with linearized p416 at the multicloning site into the *S. cerevisiae* BY4741 (MATa his3Δ1 leu2Δ0 met15Δ0 ura3Δ0) strain and selected for on SD (-) URA media. The plasmids were then extracted and transformed *en mass* into *E. coli* DH5α competent cells and plated on LB plates with 100 μg/ml ampicillin. *E.coli* colonies were miniprepped and sequenced (Plasmidsaurus Inc) to determine which were correct. The resulting correct plasmids were individually transformed into aforementioned *S. cerevisiae* strain RP4400 as described above.

### Sample Preparation of *S. cerevisiae* for Mass Spectrometry Analyses

Yeast strains were grown overnight at 30 °C in 5 mL of SD-URA medium with 2% galactose. The following morning, cultures were pelleted by centrifugation for 3 minutes at 3000 rpm. The pellet was then washed with 12 mL of autoclaved Milli-Q water, followed by centrifugation under the aforementioned conditions. The cell pellets were then resuspended with 5mL of autoclaved Milli-Q water, and the optical density (OD600) of each culture is calculated using a 1:50 culture dilution in a cuvette. The volume is then adjusted to reach a final OD600 of 5. Following this, 5 µl of drug in solvent or solvent-only is added to 495 µL of the culture for each sample. Samples were then incubated at 30 °C for 4 hours in a rotary shaker. Following incubation, samples were pulse-spun and transferred to 96-well filter plates (Pall P/N 8684) and the buffer was removed via vacuum filtration into a collection plate. Incubation media collection plates were then spun at 1000 rpm for 1 minute to remove residual droplets on the side walls. Cells were then resuspended in 50 µL of autoclaved Milli-Q water and immediately frozen at -80 °C in Safe-Lock Eppendorf tubes, whereas the incubation buffer was transferred to Safe-lock Eppendorf tubes and frozen.

Frozen yeast pellets were thawed and resuspended in 800 µL of a 50:50 mixture of BuOH and MeOH alongside 100 µL of 0.5 mm glass beads. Samples were lysed using a bead-beating device as described above, following by centrifugation at 14,000 rpm for 10 minutes. From each sample, 800 µL of the BuOH:MeOH extract was transferred into an Eppendorf Safe-Lock microtube for subsequent drying.

To dry the samples, the incubation buffer is thawed, and tubes are placed inside an Eppendorf vacufuge ensuring balance after the device has been preheated for 15 minutes at 60 °C. Samples are dried for approximately 2 to 3 hours at 60 °C (V-AL setting). When samples are dried, samples are stored at -20 °C until sample resuspension. Samples are resuspended as described above for the worm samples.

### Liquid Chromatography–Quadrupole Time-Of-Flight Mass Spectrometry

Agilent 1260 Infinity II with 6545 LC-MS/QTOF mass spectrometer was used to analyze samples in positive ionization mode with Dual AJS electrospray ionization (ESI) equipped with Agilent ZORBAX Eclipse Plus C18 column (2.1x50mm, 1.8-μm particles) and ZORBAX Eclipse Plus C18 guard column (2.1x5mm, 1.8-μm particles). LC parameters used were: injection volume 5 μL with 5 μL needle wash with sample prior to injection. The autosampler chamber temperature is maintained at 4°C and the column oven temperature is maintained at 40°C. Mass spectrometry parameters were: gas temperature 320°C, drying gas flow 8 L/min, nebulizer 35 psi, sheath gas 350°C at 11 liters per minute, VCap 3500 V, Nozzle voltage 1000 V, fragmentor 125 V, skimmer 65 V. The solvent gradient with the flow of 0.5 ml per minute started with 95% mobile phase A (Optima LC/MS H2O+0.1% Formic Acid, Fisher Chemical P/N LS118-4) and 5% mobile phase B (Optima LC/MS Acetonitrile+0.1% Formic Acid, Fisher Chemical P/N LS120-4), increased linearly to 50% B at 10 minutes, followed by an increase to 100%B at 27.50 minutes. The post-run time was two minutes (instrument conditioning at 95% mobile phase A). The raw data was analyzed using Agilent MassHunter Qualitative Analysis 12.0 and MassHunter Profinder 10.0 to find differences in samples of different strains and conditions.

### Computational Modeling

As experimentally resolved structures are not available for the complexes of interest, Boltz-1 (Wohlwend *et al*. 2025), an open source implementation of AlphaFold 3 (Abramson *et al*. 2024), was used to co-fold the enzyme, HEME, and CAD. Thirty structures were generated and the top ranked (by confidence score) model was selected for molecular dynamics simulation. All confidence, pTM, ligand ipTM, and complex pLDDT scores were better than 0.92. For simulation with AMBER (Pearlman *et al*. 1995), HEME parameters for a ferric high-spin state were adapted from (Shahrokh *et al*. 2012), and a covalent linkage was made between HEME and its coordinating cysteine, C444.

CADs were parameterized using antechamber and GAFF (Wang *et al*. 2004). The uncharged version of each CAD was setup using the identical pose as the charged version, but with a different protonation state. The systems were then solvated and neutralized with sodium ions using tleap. Initial equilibration consisted of two minimization steps (the first with protein restraints), followed by 1 ns of restrained NVT dynamics (heating from 0 K to 300 K), and 1 ns of unrestrained NPT dynamics at 300 K and 1 atm. All simulations used the AMBER ff15ipq force field (Debiec *et al*. 2016) and TIP3P water model, selected for its consistency with NMR-derived structural data (Koes AND Vries 2017). Three independent 100-ns production runs were performed for each system. Analysis was performed using MDAnalysis (Michaud-Agrawal *et al*. 2011).

Representative ligand poses for each simulation (i.e., Figure 6) were selected by identifying the medoid of the largest cluster, where clusters were determined using root mean square deviation (RMSD) as a distance metric for the fcluster method of scipy with a radius of 2.0 angstroms and average linkage.

### CAD Sensitivity Analysis

Worms were incubated with CADs in NGM liquid media supplemented with *E. coli* HB101 (OD600 ≥ 1.2) in a shaking incubator (200rpm) at 20°C for 3 days prior to quantification. Twenty worms were deposited per well (two wells per condition and genotype) in a total volume of 50 µL, where live moving worms were quantified up to a ceiling of 50. Data were normalized to each strain’s respective solvent control.

### Analysis of Gut Granules using cDCFDA

Synchronized L1 animals were grown in NGM liquid media supplemented with *E. coli* HB101 (OD600 ≥ 1.2) for 24 hours at 20°C on a tube rotator prior to addition of CADs, which were then added at the indicated final concentrations and incubated for another 24 hours. Following this, 5-(and-6)-carboxy-2’,7’-dichlorofluorescein diacetate (cDCFDA) (Cayman Chemicals Item #34692) was added to a final concentration of 50 µM and incubated for 2.5 hours. After the incubation, worms were washed seven times with M9 media, anesthetized with 250 mM sodium azide, and mounted on agarose pads. Imaging was performed using a Leica DM6B microscope with a 20x objective. Puncta were quantified using ImageJ. The GFP channel images capturing cDCF signal were processed with a Difference of Gaussians filter (σ_1_ = 1.0, σ_2_ = 1.5) to isolate punctate structures.

Images were then thresholded using the RenyiEntropy method and segmented using the Watershed algorithm. Particles were then subsequently quantified in the intestine using size (0.11-40 µM^2^) and circularity (0.45-1.00) parameters.

### The Screening and Characterization of Actinomyces Bacterial Strains for CAD Reporter Induction

65 Cubist Actinomyces strains were plated in the wells of 12 well plate with solid MYM agar at 30°C for seven days in duplicate. ∼1000 synchronized L1 RP3507 *trIs114* [*pgp-13p*::YFP; p382(*myo-2p*::mCherry)] CAD reporter worms were deposited into each well. 24 hours later wells were inspected for worms expressing YFP using an MZFLIII Leica epifluorescent dissection microscope. Subsequent detailed analysis was done by growing ‘hit’ strains plus a non-hit control on solid 6 cm MYM agar plates at 30°C for three days. 500 synchronized L1 RP3507 animals were then transferred to the bacterial lawns and incubated for 24 hours. Following co-incubation, worms were manually picked, anesthetized using 250 mM sodium azide, and mounted on agarose pads.

Imaging of reporter signals in the head of animals was performed on a Leica DM6B microscope with a 63x objective, and quantified using ImageJ.

## Acknowledgements

We thank Igor Stagljar and Jamie Snider for training and use of their CLARIOstar fluorimeter. We thank June Tan, Michael Schertzberg and Andrew Fraser for performing Linux-based calculations for the structural similarity map and guidance on sequence data analyses. We thank Jessica Lacoste, Jessica Knox, and Doreen Fang for technical assistance and the Charlie Boone and Brenda Andrews labs for the gift of the Y15771 *S. cerevisiae* yeast strain. Some *C. elegans* strains were provided by the CGC, which is funded by NIH Office of Research Infrastructure Programs (P40 OD010440). We thank Shohei Mitani and the National Bioresource Project of Japan for multiple *C. elegans* mutant strains.

## Funding

Funders include the Canadian Institutes of Health Research Project Grants 173448 and 186156 (PJR), a Canada Research Chair (Tier 1) grant in Chemical Biology (PJR), an Ontario Graduate Scholarship to LT, a grant from the National Institute of General Medical Sciences R35GM140753 (DK), and a NSERC grant RGPIN-2020-07212 (CC).

## Source Data File 1 (available upon request)

**Tab 1: Spectrum Library Data Associated with Multiple Figures.** Key columns include column J, the node identification numbers from Figure 3A; column Z, a sortable column that places the molecules capable of suppressing *pgp-14(0)* at the top with preliminary crystal counts in column AA associated with Figures 1A and 1B; columns FY and FZ, which indicate whether a given molecule is a secondary or tertiary amine; column HA which indicates whether a compound is predicted to be a phospholipidosis-inducing CAD, which is associated with Figure 3B; column HU, which indicates the 112 gold standard phospholipidosis-inducing CADs associated with Figure 3B and 3C; columns IE-IG, which shows the mini-library analysis of compounds and whether they induce pgp-13 reporter expression in a wild type or nhr-70(0) or nhr-107(0) mutant background, associated with Figure 4.

**Tab 2: CAD and wact-190 Data Associated with Figure 1C**. Column A and B list the 14 distinct CAD treatments given at 30 µM. Columns C-H represents the average viability of technical replicates for wild-type animals treated with solvent or drug alone. Columns I-N represent the viability of wild type animals treated with wact-190 (30 µM) in the absence or presence of each of the 14 CADs. Columns O-T represents the viability of *pgp-14(ok2660)* animals treated with solvent or drug alone. Columns U-Z represents the viability of *pgp-14(ok2660)* animals treated with wact-190 in the absence or presence of each of the 14 CADs. Column AC contains the statistical analyses using a two-tailed Student’s t-test, comparing the *pgp-14(ok2660)* worms treated with CADs in the presence of wact-190 to those treated with wact-190 alone. This data supports Figure 1C.

**Tab 3: Crystal Formation Data Associated with Figure 1D**. Column A and B list the 14 distinct CAD treatments given at 30 µM. Columns C-I represents the percentage of wild-type animals with observed crystals when treated with the CADs alone. Columns J-P demonstrates the percentage of wild-type animals with crystals when given wact-190 (30 µM) in the absence or presence of each of the 14 CADs. Columns Q-W represents the percentage of *pgp-14(ok2660)* animals with crystals when treated with each of the 14 CADs alone. Columns X-AD represents the percentage of *pgp-14(ok2660)* animals with observed crystals when treated with wact-190 in the absence or presence of each of the 14 CADs. Column AF contains the statistical analyses using a one-tailed t-test comparing crystal re-sensitization in *pgp-14(ok2660)* animals treated with CADs in the presence of wact-190 compared to those treated with wact-190 alone. This data supports Figure 1D.

**Tab 4: Forward Genetic Screen Sequencing Analysis Data Associated with Figure 2A**. Column A represents the mutation call number; column B, the chromosome number where the mutation was detected; columns C and D indicate the start and end positions of the mutation; Column E is the reference base, and column F is the base read in the mutants; Column G indicates the genomic region where the mutation occurred; Column H is the annotated gene associated with the mutation, whereas column L indicates the specific type of mutation if the variant is exonic. Columns T-BJ represents each of the sequenced mutants and indicates whether a homozygous or heterozygous mutation was detected.

**Tab 5: Mutant Dose-Response Data Associated with Figure 2A**. Key columns include columns A-S, which represents the raw data for three biological replicates of wild-type, unmutagenized (parental) RP3412, and the mutant animals from the forward genetic screen treated with a range of concentration of the CAD terconazole, both alone and in combination with wact-190 at (30 µM). Columns AC-AH contains the averages calculated from each replicate, which helped calculate the IC50 in Figure 2A.

**Tab 6: Mutant Dose-Response Data in Combination with wact-190 Associated with Figure 2A**. Key columns include column B, which denotes the strain, and column C, which denotes the genotype. Column D-K indicates the average viability from each of the biological replicates at each concentration of terconazole combined with 30 µM of wact-190. Column N represents the IC50 based on the preceding columns, and column O represents the average percentage of animals with observed crystals when given the combination of 30 µM Terconazole and 30 µM wact-190.

**Tab 7: Mutant Crystal Formation Data Associated with Figure 2A**. Columns A-K reflect the raw counts of crystal observation for each strain with each treatment – either solvent, 30 µM terconazole, 30 µM wact-190, or a combination of 30 µM terconazole and 30 µM wact-190. Column T has each strain with each treatment noted, whereas column U through W represents the percentage of animals with crystals from each biological replicate, and column Y indicates the mean +/- the standard error for each condition.

**Tab 8: qRT-PCR Gene Expression Data Associated with Figure 2C**. Key columns include column A indicating the genotype, whereas Column B is the strain used. Columns E-P represent the average relative transcript expression levels of specific genes – *pgp-14* (columns E-F), *pgp-13* (columns G-H), *cyp-35b1* (columns I-J), *cyp-35b2* (columns K-L), *cyp-35b3* (columns M-N), and *cror-1* (columns O-P) – with animals treated with either solvent (DMSO) or 30 µM terconazole.

**Tab 9: *pgp-13p*::YFP Reporter Time-Course Data Associated with Figure 2H-2I**. Key columns include column C, which indicates the time point that fluorescence measurements were taken. Columns E-G represents the worms given solvent only, whereas columns I-K represents worms given 15 µM terconazole or amiodarone, and columns M-O represents the YFP recorded from worms given 30 µM terconazole or amiodarone. Columns Q and R represent the two-tailed t-test statistical analyses.

**Tab 10: *pgp-13p*::YFP Reporter Dose Response Data +/- NHR-70 and NHR-107 Associated with Figure 2J-2K**. Columns A-F denotes dose-response data for each strain –wild type, *nhr-70(0),* and *nhr-107(0)* – animals given Terconazole at a range of concentrations with YFP measurements. Column H-M reports similar data, with the difference being worms are given Amiodarone. Columns A and H indicate the dose, Column C and E indicate the measurements from each trial, and Columns F and M indicate the average.

**Tab 11: Evaluation of Different Methods of Calculating LogP on the PLD-CAD Prediction Model Associated with Figure 3**.

**Tab 12: Statistical Analyses of the Different Physicochemical Properties of the Molecules that are NHR-70 and/or NHR-107-Dependent, Associated with Figure 4B**. See Rows 2410-2418 for details.

**Tab 13: *C. elegans* LCMS Data Associated with Figure 5A-5B**. This tab contains metabolite data for worms given the CADs Azelastine, Chlorprothixene, Clomipramine, Clozapine, Eticlopride, and Terconazole. Key columns include E, AQ, EO, GX, HW, and JV - which indicates which mutant ratio is being presented. This is followed by the ratio data for each parent molecule or metabolite in columns F-AD, AR-DQ, EP-GD, GY-HK, HX-JJ, and JW-KV. One-tailed t-tests follow the metabolite data, in columns AF-AL, DS-EJ, GF-GR, HM-HQ, JL-JQ, KX-LA.

**Tab 14: *S. cerevisiae* LCMS Data Associated with Figure 5C**: This tab contains metabolite data for yeast given the CADs given Azelastine, Chlorprothixene, Clemastine, Clemizole, Clomiphene, Clomipramine, Clozapine, Eticlopride, Fipexide, Loxapine, and Terconazole. Columns C-I contains the ratio for CYP35-B1,B2, and B3 expressing yeast compared to control for the parent CAD. Columns K-Q contains the ratio for CYP35-B1,B2, and B3 expressing yeast compared to control for the mono-oxidized CAD metabolite. Columns S-Y contains the ratio for CYP35-B1,B2, and B3 expressing yeast compared to control for the di-oxidized CAD metabolite. Columns AA-AJ contains the one-tailed Student’s t-test statistical analyses.

**Tab 15: *S. cerevisiae* CYP-35B2 Variant LCMS Data Associated with Figure 6C**: This tab contains metabolite data for yeast given the compounds AB1515, chlorprothixene, and clomipramine. Columns A-W contains AB1515 associated data, Columns AE-BY contains chlorprothixene associated data, and Columns CK-DZ contains clomipramine associated data. Key columns include columns C, AG and CM which denotes which strain data is presented – including empty vector, CYP-35B2, as well as D311N and D311G variants. Columns I-W, AM-BY, and CT-DZ contains the metabolite and parent counts for AB 1515, chlorprothixene, and clomipramine, respectively.

**Tab 16: CAD Viability Data Associated with Figure 7A**: This tab contains normalized viability data for *C. elegans* given 6 distinct CADs – azelastine, clemizole, clomiphene, clomipramine, prochlorperazine, and terconazole – at a range of concentrations. Column A denotes the strain and genotype; column B denotes the CAD. Columns C-Z denotes the normalized viability data for each condition.

**Tab 17: cDCF Puncta Data:** This tab contains puncta counts for *C. elegans* given solvent or the CADs clomipramine or chlorprothixene. Key columns include column A, denoting the strain, column B denoting the drug, column C denoting the concentration, and column D denoting the biological replicate. Columns E-N represents counts from each worm, with the averages in column O. Rows 20-34 and 90-139 includes statistical analyses for the preceding data sets. This data supports Figure 7C and 7D.

**Tab 18: *Streptomyces pgp-13p*::YFP Reporter Measurements Associated with Figure 8C**: This tab contains *pgp-13p*::YFP fluorescence measurements for *C. elegans* placed on three distinct strains of *Streptomyces* – 5E1 (negative), 1G9, and 1E6 (both positive hits) for twenty-four hours. Column A denotes the strain of *C. elegans*, Column B denotes the strain of *Streptomyces*, Column C denotes the biological replicate number, and Columns D-M indicates the individual measurement from the head of the worm, and column N is the average across the ten worms.

**Tab 19: Crystal Formation Data Associated with Supplemental Figure 3B.** This tab contains data pertaining to crystal formation for wild type, RP2960, RB894, TM1697, RP3522, TM4512, RP3523 and RP3495 given either solvent, 30 µM terconazole, 30 µM wact-190, or a combination of the two drugs. Column A denotes the strain, Column B denotes the genotype and treatment, Column C-J denotes the observation counts for crystal presence, whereas column L-N denotes the calculated percentage of crystals observed. Column P denotes statistical analyses for a two-tailed Student’s t-test comparing the strains given just 30 µM wact-190 compared to those given 30 µM terconazole and 30 µM wact-190.

**Tab 20: Crystal Formation Data Demonstrates PGP-13 Required for wact-190 Re-sensitization Associated with Supplemental Figure 4B.** This tab contains data pertaining to crystal formation for wild type, RP2960, RB894, RP3495, and RP3561 given either solvent, 30 µM terconazole, 30 µM wact-190, or a combination of the two drugs. Column A denotes the strain, Column B denotes the genotype and treatment, Column C-J denotes the observation counts for crystal presence, whereas column L-N denotes the calculated percentage of crystals observed. Column P denotes statistical analyses for a two-tailed Student’s t-test comparing the strains given just 30 µM wact-190 compared to those given 30 µM terconazole and 30 µM wact-190.

**Tab 21: qRT-PCR Data for *wt*, *nhr-70(0),* and *nhr-107(0)* given 5 CADs Associated with Supplemental Figure 5.** This tab contains qRT-PCR based gene expression data pertaining to *wt*, *nhr-70(0)*, and *nhr-107(0)* worms given 5 CADs – namely terconazole, clomipramine, azelastine, clemizole, and clomiphene. Column A denotes the strain and treatment, Column B denotes the transcript being measured, and Columns C-E denotes the average fold change across three trials. Columns H-M and O-T contains the statistical analyses comparing treatments within strains, and across strains within the same treatment, respectively.

**Tab 22: The Analysis of the Accumulation of the CAD Parent Molecules in Yeast Cells as Measured by LCMS, Associated with Supplemental Figure 6.**

---

**Supplemental Figure 1:**
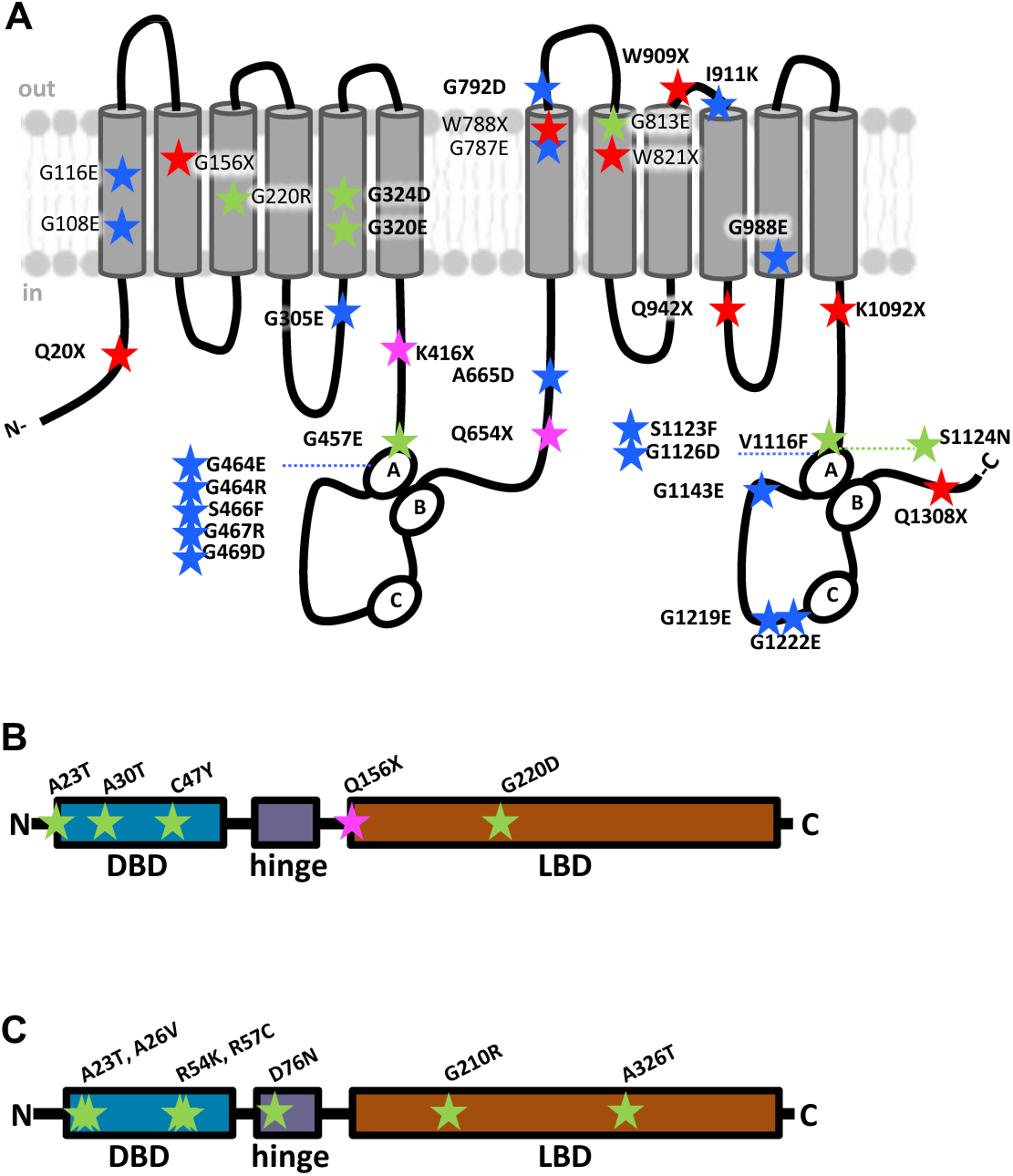
A Schematic of Mutations Recovered in our Forward Genetic Screen. **A.** A topology schematic of PGP-13 and mutations identified in our screen with missense in green and nonsense in pink. Previously recovered PGP-14 missense (blue) and nonsense (red) mutations are also shown. **B.** Topology schematic of NHR-70 nuclear hormone receptor and associated mutations. **C.** Topology schematic of NHR-107 nuclear hormone receptor and associated mutations. In B & C, green stars indicate a missense and pink star indicate nonsense.

**Supplemental Figure 2.**
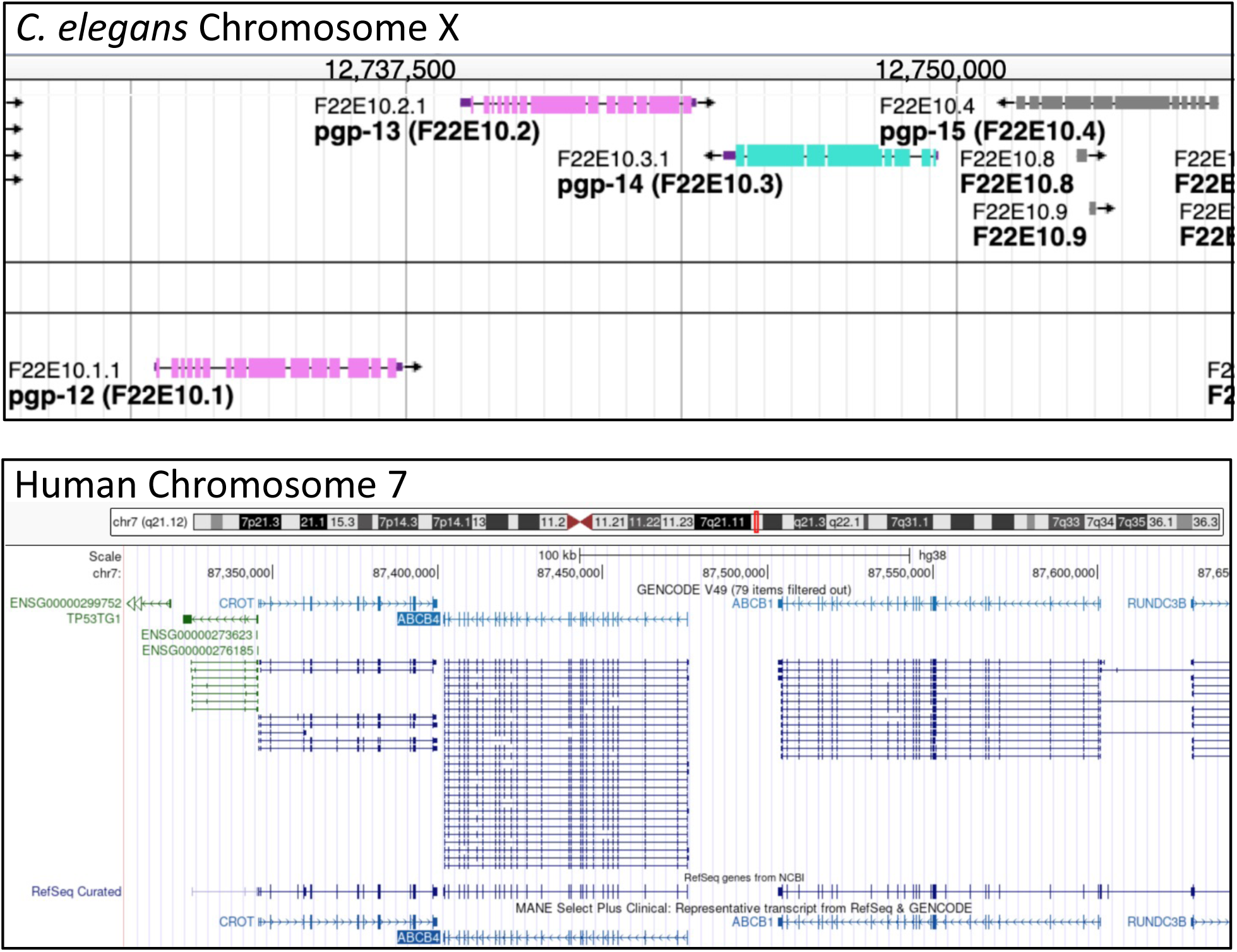
A Comparison of the Relative Positions of Relevant PGPs in *C. elegans* and Humans. The screenshot of the genomic position of *pgp-13* and *pgp-14* on chromosome X is from WormBase (version WS297). The screenshot of the genomic position of ABCB4 and *ABCB1* on chromosome 7 is from the UCSC Genome Browser (accessed Nov 12, 2025).

**Supplemental Figure 3.**
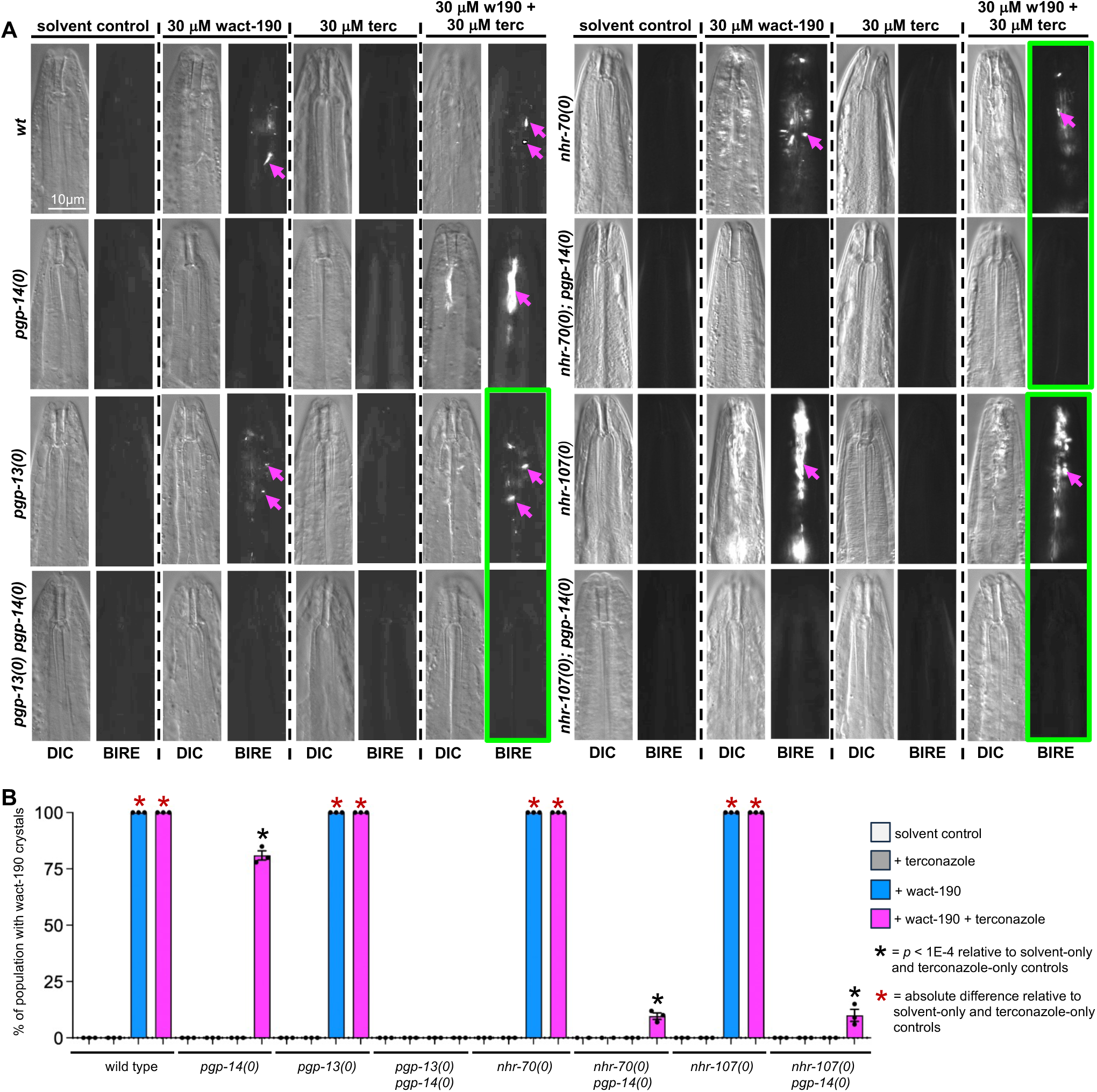
PGP-13, NHR-70, and NHR-107 are Necessary for CAD Resensitization of *pgp-14(0)* to wact-190 Crystal Formation. **A.** Images of the heads of animals of the indicated genotype (to the left) and treatment (at the top of the columns). Treatment time was 48 hours. The imaging channel for each animal is indicated at the bottom of the columns; DIC, differential interference contrast; BIRE, birefringence. Anterior is up. The scale in the top left applies to all images. The relevant strains/genotypes are N2 *wild type*; RP2960 *pgp-14(ok2660)*; RB894 *pgp-13(ok747);* RP3495 *pgp-13(tr696) pgp-14(ok2660)*; *nhr-70(tm1697),* RP3522 *nhr-107(tm1697); pgp-14(ok2660)*; *nhr-107(tm4512)*; and RP3523 *nhr-107(tm4512); pgp-14(ok2660)*. wact-190 crystals are indicated with a pink arrow. The key result that demonstrates each of PGP-13, NHR-70, and NHR-107 are necessary for the CAD (terconazozle) to resensitize the *pgp-14(0)* null to wact-190 crystal formation are indicated with green box. **B.** Quantification of the number of animals within the population that exhibited wact-190 crystals in the experiment shown in A. N=3 independent trials with at least 24 animals counted per sample (n≥24). A two-tailed Student’s T-test was used to determine *p* values, except where there are absolute differences (all trials for a sample yields zero or 100 percent).

**Supplemental Figure 4.**
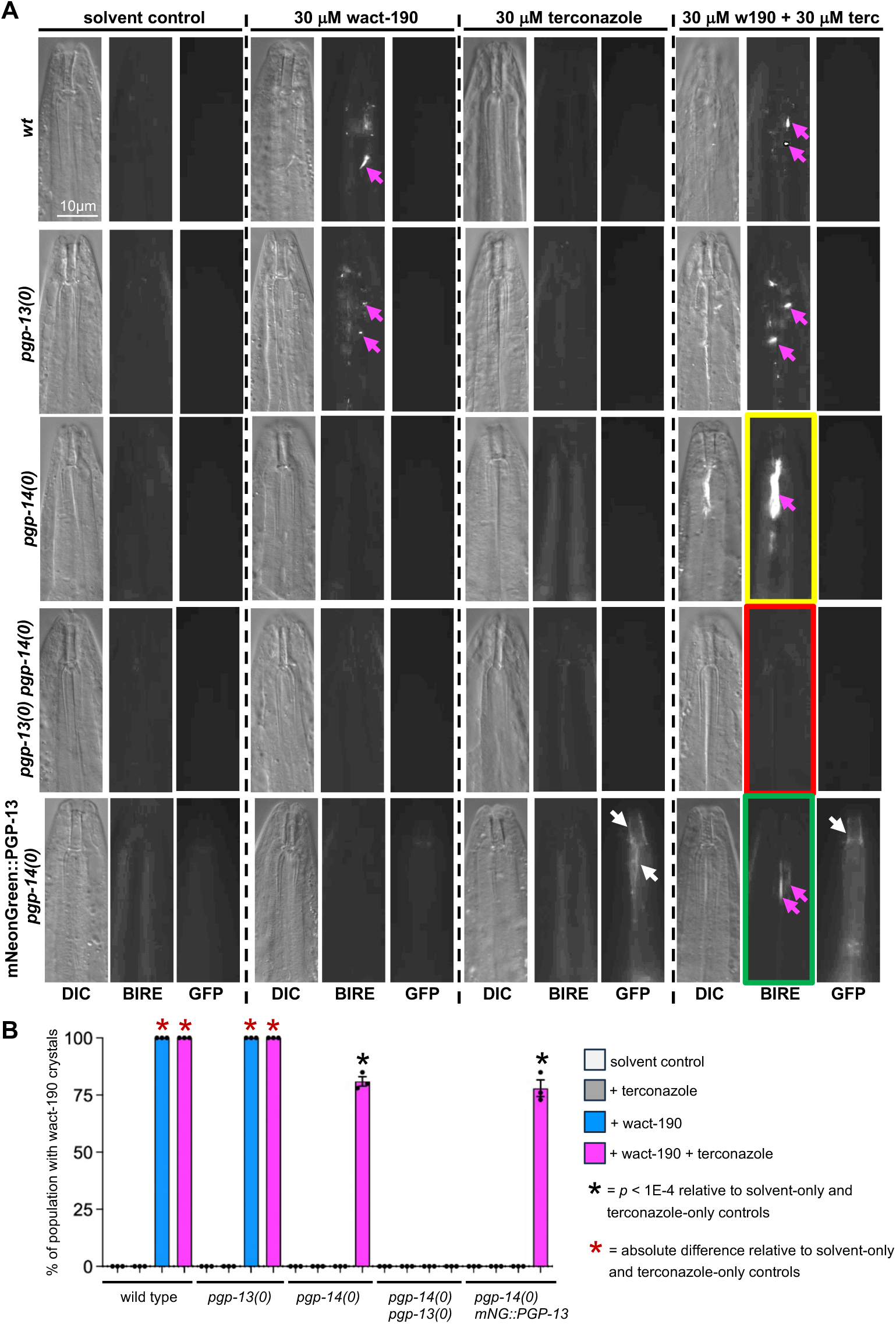
The mNeonGreen::PGP-13 Translational Fusion is Functional. **A.** Images of the heads of animals of the indicated genotype (to the left) and treatment (at the top of the columns). Treatment time was 48 hours. The imaging channel for each animal is indicated at the bottom of the columns; DIC, differential interference contrast; BIRE, birefringence; GFP, green fluorescent protein. Anterior is up. The scale in the top left applies to all images. The relevant strains/genotypes are N2 *wild type* ; RB894 *pgp-13(ok747);* RP2960 *pgp-14(ok2660)*; RP3495 *pgp-13(tr696) pgp-14(ok2660)*; RP3561 *trIs115 [mNeonGreen::PGP-13] pgp-14(ok2660)*. wact-190 crystals are indicated with a pink arrow; mNeonGreen::PGP-13 signal is indicated with white arrows. The key result that demonstrates mNeonGreen::PGP-13 functionality is that the strain containing the mNeonGreen insertion into PGP-13 in the background of the *pgp-14(ok2660)* null yields crystals (indicated by a green box) in the background of the CAD terconazole (which suppresses *pgp-14(0)* because it upregulates *pgp-13* expression), just like wild type *pgp-13* in the background of *pgp-14(0)* (yellow box). If *pgp-13* loses its function, terconazole cannot resensitize *pgp-14(0)* to wact-190 crystals (red box). Note that the controls for this experiment are the same as that shown in Supplemental Figure 3 (the experiments were done at the same time). **B.** Quantification of the number of animals within the population that exhibited wact-190 crystals in the experiment shown in A. N= 3 independent trials with at least 24 animals counted per sample (n≥24). A two-tailed Student’s T-test was used to determine *p* values, except where there are absolute differences (all trials for a sample yields zero or 100 percent).

**Supplemental Figure 5.**
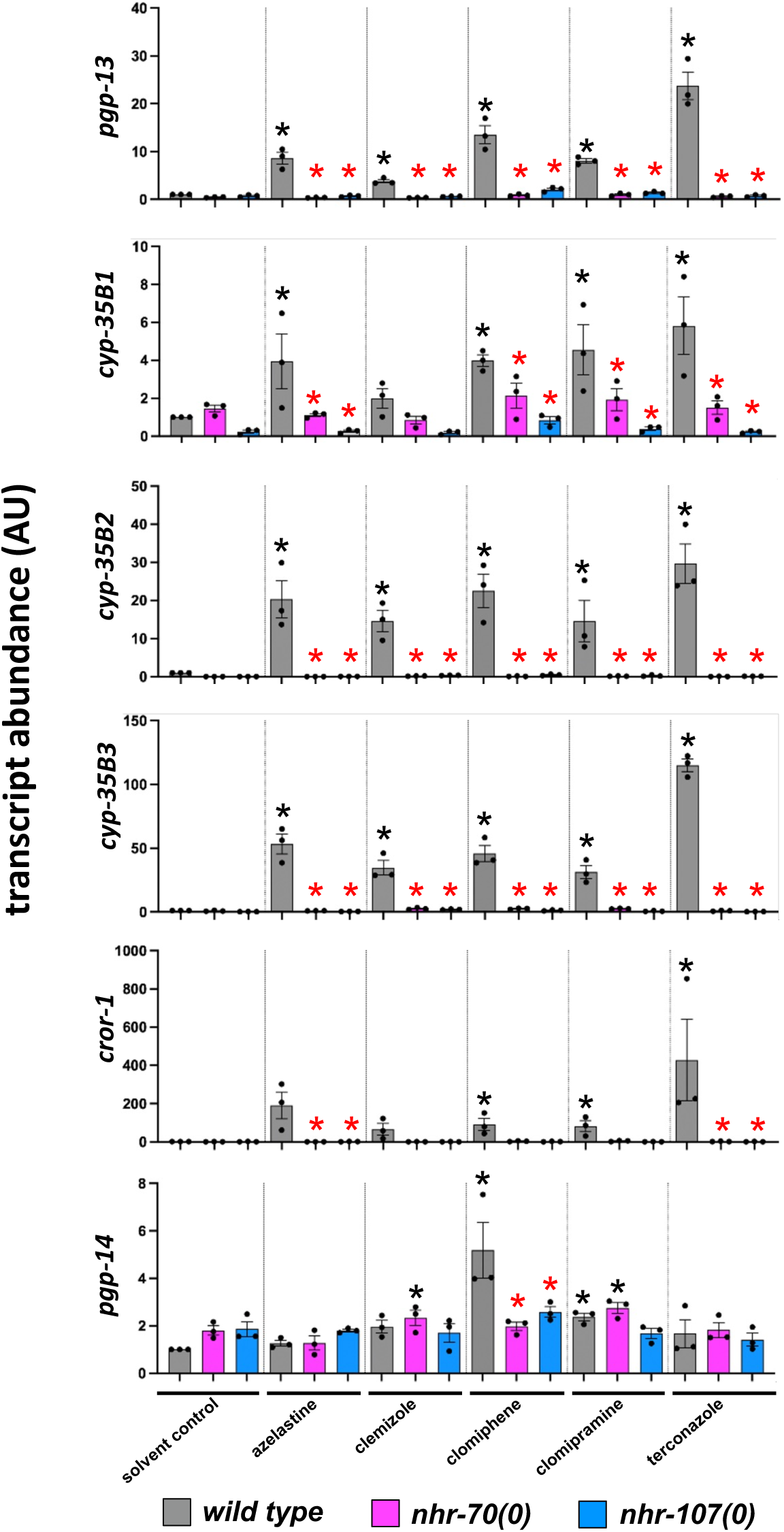
The qRT-PCR analysis of *pgp-13*, *pgp-14, cyp-35b1*, *cyp-35b2*, and *cyp-35b3* transcript abundance in *wild type*, *nhr-70(tm1697)*, and *nhr-107(tm4512)* animals. The worms were treated for 96 hours with either solvent control, terconazole (30 µM), clomipramine (30 µM) clomiphene (20 µM), azelastine (20 µM), or clemizole (20 µM). Data shown represents mean ± S.E.M.; N=3 biological trials with n=2 technical repeats each. Statistical significance was determined via two-way ANOVA followed by Fisher’s LSD test comparing each treatment to the solvent control within each respective genotype. Black asterisks indicate statistical significance of *p*<0.05 relative to the solvent-only control. Red asterisks indicate significance of *p*<0.05 between the mutant strains compared to wild type within each treatment group.

**Supplemental Figure 6.**
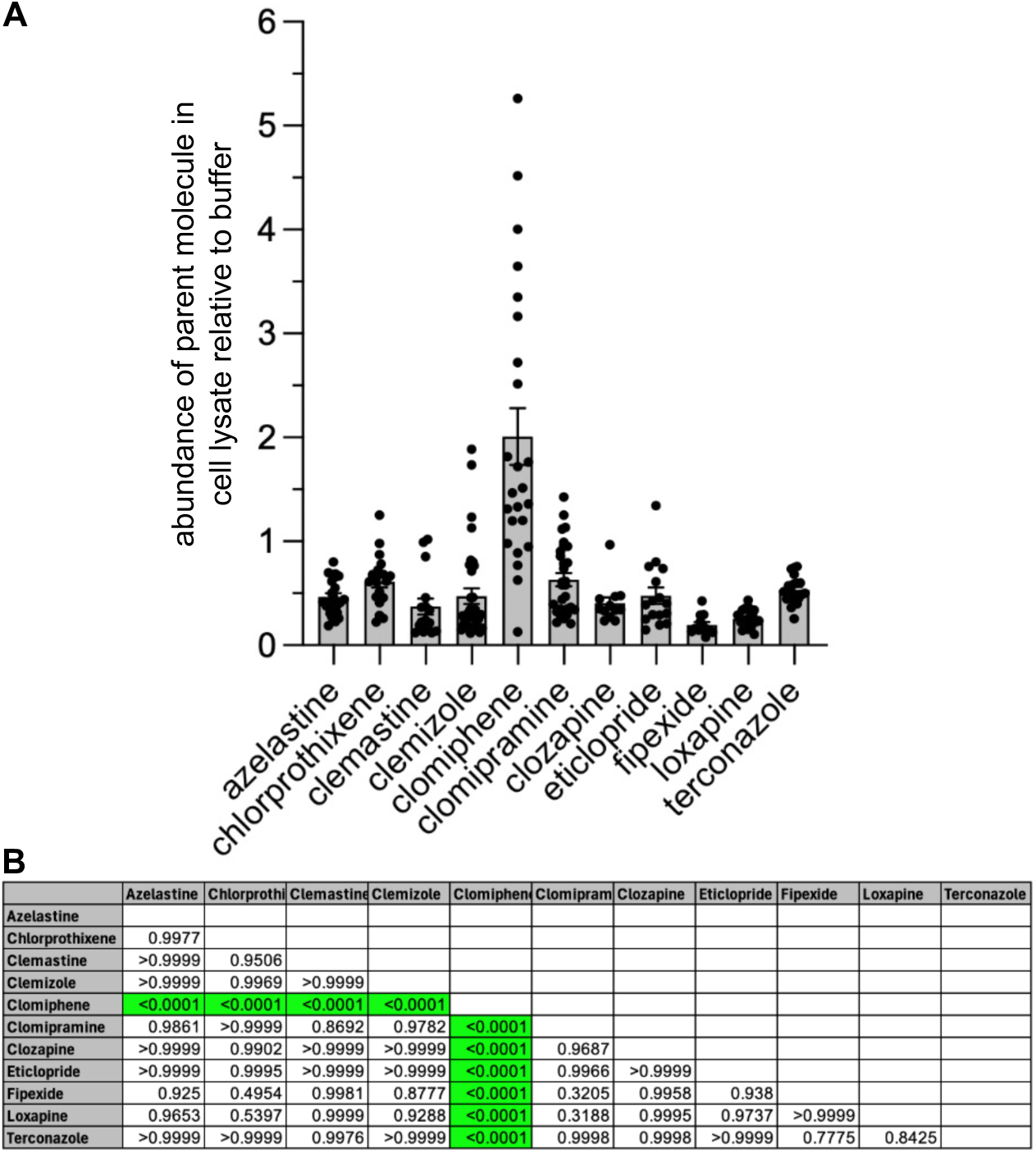
The Accumulation of CAD Parent Molecules in Yeast Cells. **A.** For every independent repeat of every experiment shown in Figure 5c (N,≥12 independent repeats), we calculated the ratio of the parent (i.e., non-metabolized) CAD molecule that is present in the cell lysate versus that found in the buffer. With the assembled dataset, we performed a one-way ANOVA analyses revealing *p*<0.001. Standard error of the mean is shown. **B.** An all-against-all statistical comparison was performed on the data shown in A using the Tukey’s multiple comparisons test. Only the accumulation of clomiphene in yeast is significantly different than other CADs (*p*<0.05).

**Supplemental Figure 7.**
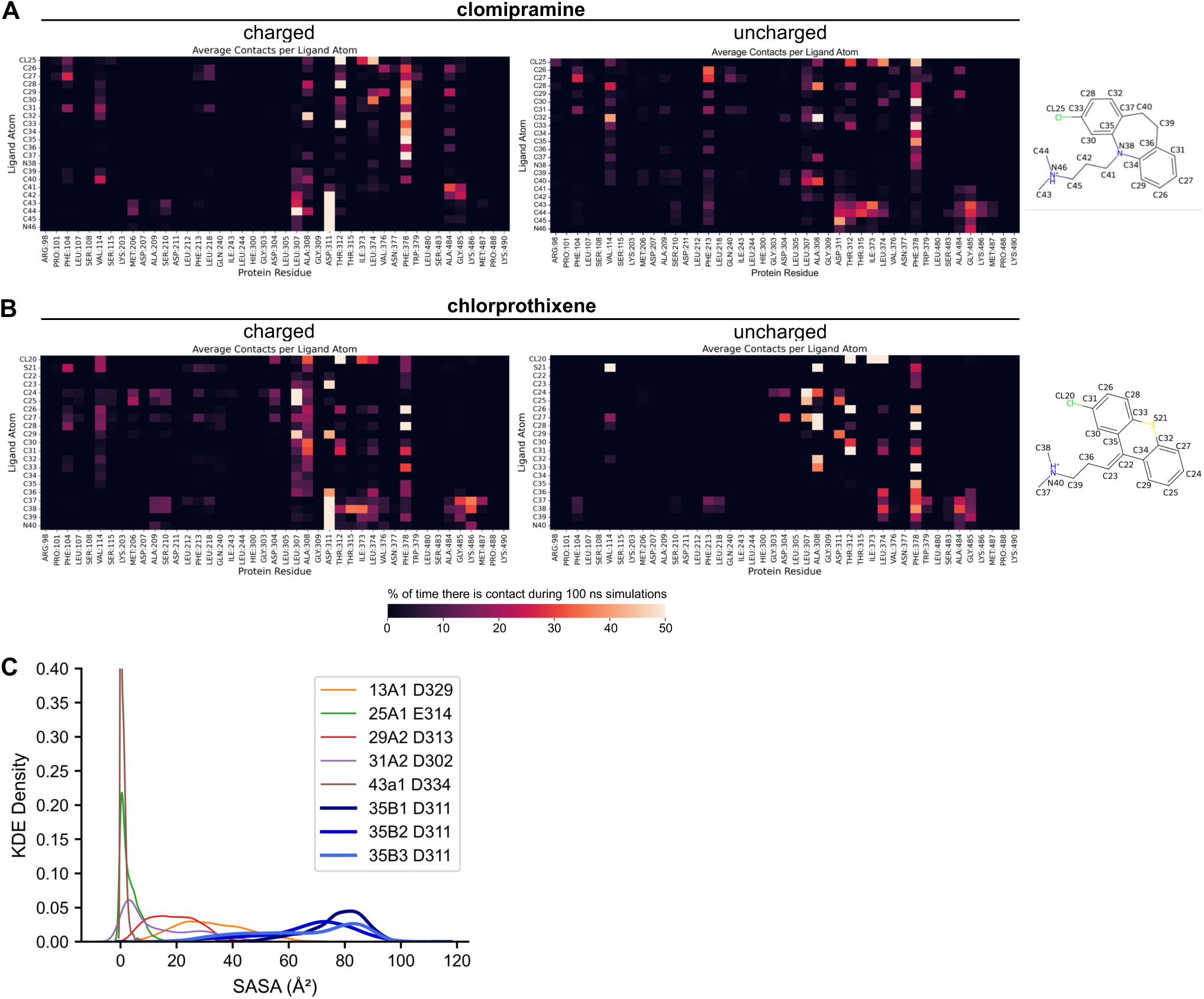
A Survey of the Potential CYP35-B2 Binding Site Residue Contacts with Exemplar CADs. **A-B.** Each heatmap for the indicated charged and uncharged CAD shows the percent of time during 3 x 100ns simulation that each atom of the CAD ligand was in contact (<4 Å) with any atom of a given CYP-35B2 binding site residue. The difference in charge is due to different protonation of the nitrogen (bottom row of all the heatmaps). The atom numbers on the left of the charts correspond to those indicated on the molecules to the right of the charts. The legend below B is for all charts. **C.** Illustration of the solvent accessible surface area of selected *C. elegans* P450s showing that even though D311 is conserved in sequence, it is not conserved in a structural role. For a selection of CYPs the apo structure was predicted with Boltz-1 and simulated for 3 x 100ns with Amber, and the solvent accessible surface area of D311 or the equivalent residues was measured. Only in CAD metabolizing P450s of the CYP-35B family was this residue highly accessible and therefore available for interacting with substrates.

**Supplemental Figure 8.**
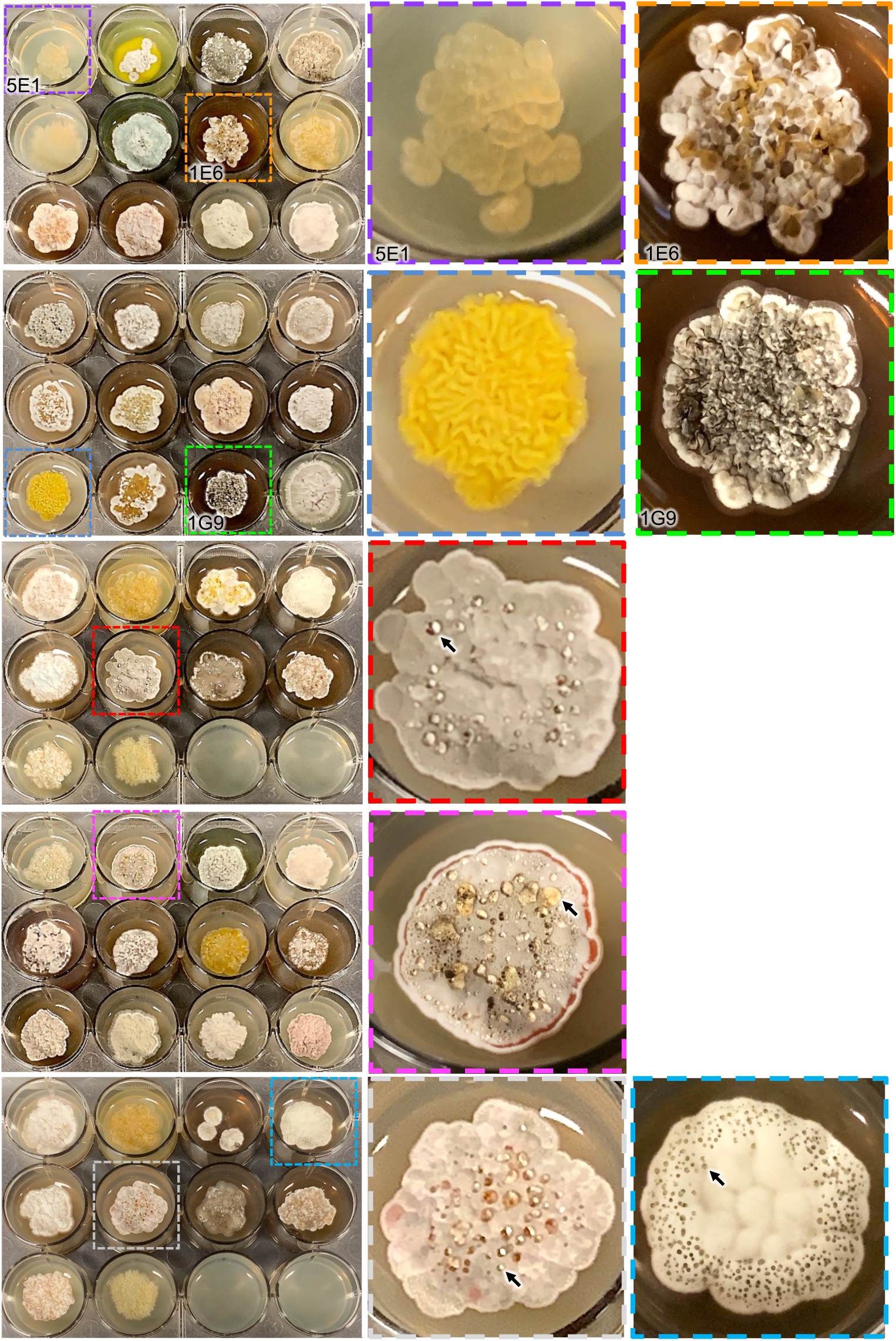
Examples of the Varied Colony Morphologies of Different *Streptomyces* Strains, Including the Hits (1E6 and 1G9) and Exemplar Non-Hit (5E1). Strains were grown in 12-well plates over the course of 7 days. In addition to the hits and non-hit, additional strains are magnified to highlight features of interest or the obvious secreted secondary metabolites (black arrows) that some strains make under these culture conditions. Also note the differences in the colour of the agar base for some strains, which also reflects the secretion of different secondary metabolites.

## Notes

### Competing Interest Statement

The authors have declared no competing interest.

